# Cell autonomous versus systemic Akt isoform deletions uncovered new roles for Akt1 and Akt2 in breast cancer

**DOI:** 10.1101/2020.01.08.898767

**Authors:** Xinyu Chen, Majd M. Ariss, Gopalkrishnan Ramakrishnan, Veronique Nogueira, Catherine Blaha, William Putzbach, Abul B. M. M. K. Islam, Maxim V. Frolov, Nissim Hay

## Abstract

Studies in three mouse models of breast cancer identified profound discrepancies between cell autonomous and systemic Akt1 or Akt2 deletion on breast cancer tumorigenesis and metastasis. First, unlike systemic Akt1 deletion, which inhibits metastasis, cell autonomous Akt1 deletion does not. Second, systemic Akt2 deletion does not inhibit mammary tumorigenesis and metastasis, but cell autonomous Akt2 deletion eliminates ErbB2 expressing cells in the mammary gland and prevents tumorigenesis. However, the elevation in insulin by Akt2 systemic deletion hyperactivates tumor Akt, enabling ErbB2 expression, and exacerbates mammary tumorigenesis. Decreasing insulin level inhibits accelerated tumorigenesis by systemic Akt2 deletion. Single cell mRNA sequencing revealed that systemic Akt1 deletion maintains the pro-metastatic cluster within primary tumors but ablates pro-metastatic neutrophils. Systemic Akt1 deletion inhibits metastasis by impairing the survival and mobilization of tumor-associated neutrophils. Importantly, neutrophil-specific deletion of Akt1 is sufficient to exert resistance to metastasis. The results underscore the importance of determining systemic effects rather than cell autonomous effects as a proof of concept for cancer therapy.

---

The serine/threonine kinase Akt is frequently hyperactivated in breast cancer through multiple mechanisms, including PI3K activation, PTEN loss, and ErbB2/Her2/neu activation/amplification ^1^. However, previous studies regarding the roles of Akt isoforms in breast cancer did not provide a coherent understanding, and the results are controversial. The ablation of Akt1 in cell culture increased the migration and epithelial mesenchymal transition (EMT), whereas ablation of Akt2 decreased EMT ^2^. In a mouse model of breast cancer, the activation of Akt1 increased tumor development but decreased metastasis, whereas the activation of Akt2 did not increase tumor development but increased metastasis ^3, 4^. Finally, in mouse models of breast cancer, the germ line deletion of Akt1 inhibits both tumor development and metastasis ^5, 6^, whereas the germ line deletion of Akt2 enhances both tumor development and metastasis ^5^. However, these studies do not distinguish between the cell autonomous and non-cell autonomous effects of Akt isoforms on breast cancer and do not emulate drug therapy. Therefore, we launched a comprehensive approach to examine the roles of Akt1 and Akt2 in mammary gland tumor initiation, progression and metastasis, and to understand their therapeutic implications. We employed three mouse models of breast cancer; two Her2 enriched models and one luminal B model, to distinguish between cell autonomous and systemic effects after tumor onset (Fig. S1).

## The effects of Akt1 versus Akt2 cell autonomous deletion on primary tumors and metastasis driven by ErbB2

In the cell autonomous mouse model (*MMTV-Neu-IRES-Cre (NIC)*), where ErbB2 and Cre recombinase are concomitantly expressed (Fig. 1a), we found that the cell autonomous deletion of Akt1 impaired tumor development and increased tumor-free survival (Fig. 1b), which is consistent with the effects of the germline deletion of Akt1 ^5, 6^. Surprisingly, the cell autonomous deletion of Akt2 completely inhibited tumor development (Fig. 1b). The heterozygous deletion of Akt2 did not inhibit tumor development (Fig. 1b), indicating that the complete inhibition of tumor development by Akt2 deletion in the mammary gland is not due to transgene (ErbB2) silencing. This finding is in stark contrast to effects of the germ line or systemic deletion of Akt2, which did not inhibit tumor development, but rather exacerbated tumor development ^5^ (Fig. 2). Immunoblot analysis showed that Akt1 was deleted in ErbB2-expressing mammary glands of *MMTV-NIC;Akt1^f/f^* mice, but no expression of ErbB2 or deletion of Akt2 was found in the mammary glands of *MMTV-NIC;Akt2^f/f^* mice (Fig. 1c). Taken together, these results suggest that ErbB2 expression cannot be tolerated in the absence of Akt2 and that cells expressing ErbB2 in the absence of Akt2 are eliminated. The results also suggest that the germ line ^5^ or systemic deletion of Akt2 (Fig. 2) enables ErbB2/Akt2^-/-^ mammary gland tumor cells to proliferate and overcome the intolerance to ErbB2 expression in the absence of Akt2. To further verify these possibilities, we generated *MMTV-NIC;R26Luc^LSL^* and *MMTV-NIC;Akt2^f/f^;R26Luc^LSL^* mice, in which the luciferase (Luc) gene was inserted in the ubiquitously expressed Rosa26 (R26) locus and is expressed only if Cre recombinase is also expressed ^7^ (Fig. 1d). If Cre recombinase is expressed and the cells survive, luciferase will be expressed and detected in the mice by luminescence imaging. Our results showed luciferase expression only in the *MMTV-NIC;R26Luc^LSL^*, *MMTV-NIC;Akt1^f/f^;R26Luc^LSL^*, and *MMTV-NIC;Akt2^+/f^;R26Luc^LSL^* mice but not in the *MMTV-NIC;Akt2^f/f^;R26Luc^LSL^* mice (Fig. 1d). These results further established the notion that the expression of ErbB2 in the absence of Akt2 is not tolerated in mammary gland cells; thus, these cells are eliminated. It remains to be explained, however, why ErbB2 expression in the absence of Akt1 can be tolerated. One potential explanation is that ErbB2 expression cannot be tolerated when total Akt activity is reduced below a certain threshold level. Akt2 is expressed at the highest level and Akt1 is expressed at the lowest level at early stages of tumor development (Fig. 1e). Therefore, it is possible that the deletion of Akt2 reduces total Akt activity more than the deletion of Akt1 at early stages of tumor development. Further support for this assertion is shown and discussed below (Fig. 2h, 2i, and supp. Fig. 3c).

**Figure 1:**
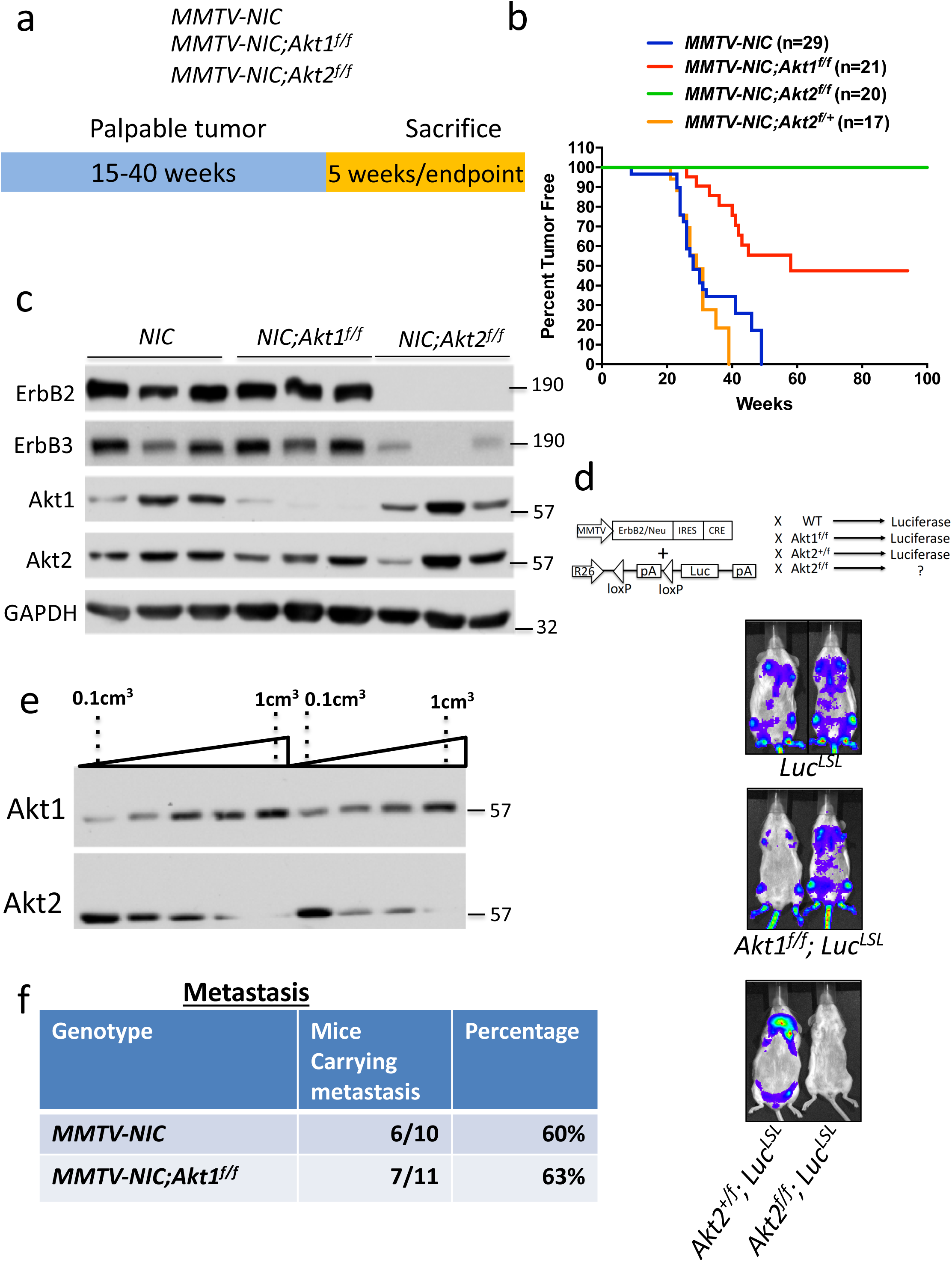
The effect of the Akt1 or Akt2 cell autonomous deletion on ErbB2-mediated mammary primary tumors and metastasis. **a.** Schematic showing the Neu-IRES-Cre (NIC) transgene and breeding scheme to generate *MMTV-NIC-Akt1^f/f^* and *MMTV-NIC-Akt2^f/f^* mice. **b.** Kaplan-Meier plots showing tumor-free survival of *MMTV-NIC*, *MMTV-NIC-Akt1^f/f^*, *MMTV-NIC-Akt2^f/f^* and *MMTV-NIC-Akt2^+/f^* mice. The number of mice is indicated. p<0.0001 *MMTV-NIC* versus *MMTV-NIC-Akt1^f/f^* and *MMTV-NIC-Akt2^f/f^*, p>0.05, *MMTV-NIC* versus *MMTV-NIC-Akt1^+/f^* using the log-rank test. **c.** Immunoblot showing ErbB2, ErbB3, Akt1 and Akt2 protein expression in the mammary glands of *MMTV-NIC*, *MMTV-NIC;Akt1^f/f^* and *MMTV-NIC;Akt2^f/f^* mice. **d.** Luminescent imaging showing luciferase expression in *MMTV-NIC;LSL-Luc*, *MMTV-NIC;Akt1^f/f^;LSL-Luc, MMTV-NIC;Akt2^+/f^;LSL-Luc*, and *MMTV-NIC;Akt2^f/f^;LSL-Luc* mice. **e**. Representative immunoblot showing the expression of Akt1 and Akt2 during primary tumor development in *MMTV-NIC* mice. **f.** Table summarizing the incidence of lung metastasis in *MMTV-NIC* and *MMTV-NIC;Akt1^f/f^* mice. The mice were sacrificed at the primary tumor endpoint, and the lungs were scored for metastasis.

**Figure 2:**
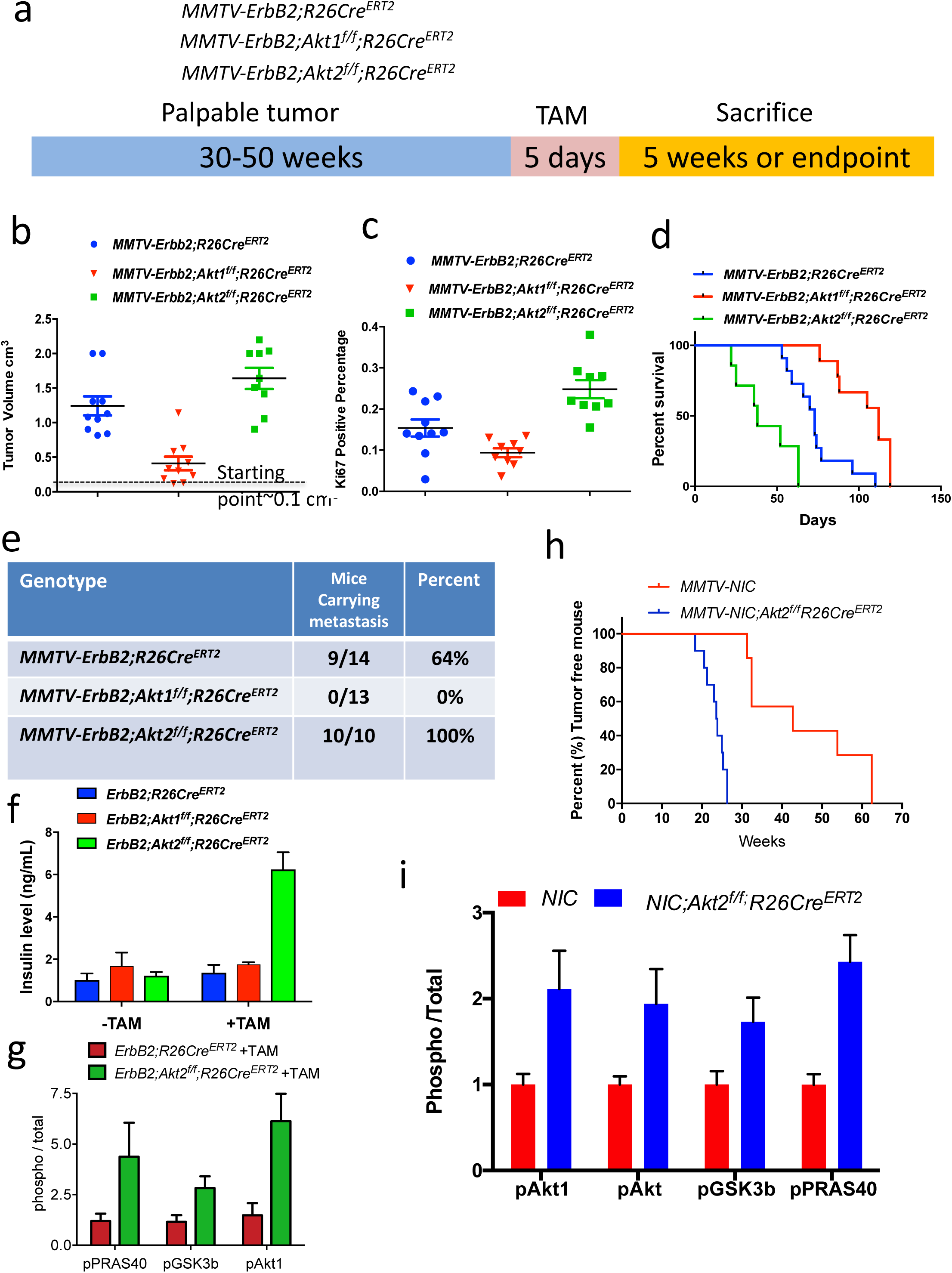
Consequences of the systemic deletion of Akt1 or Akt2 after tumor onset in *MMTV-ErbB2* mice. **a.** The genotypes of mice and experimental strategy. **b.** Tumor volume at 5 weeks after tamoxifen injection. Data are presented as the means ±SEM. p<0.001 *MMTV-ErbB2;Akt1^f/f^;R26Cre^ERT2^* vs. *MMTV-ErbB2;R26Cre^ERT2^*, p<0.025 *MMTV-ErbB2;Akt2^f/f^;R26Cre^ERT2^* vs. *MMTV-ErbB2;R26Cre^ERT2^,* using an unpaired t test. **c.** Percentage of Ki67-positive cells in tumor sections. Data are presented as the means ± SEM. P=0.017, *MMTV-ErbB2;Akt1^f/f^;R26Cre^ERT2^* vs. *MMTV-ErbB2;R26Cre^ER^.* P=0.006, *MMTV-ErbB2;Akt2^f/f^;R26Cre^ERT2^* vs. *MMTV-ErbB2;R26Cre^ERT2^* using an unpaired t test. **d.** Kaplan-Meier survival curves as determined by tumor endpoint. P=0.0007, *MMTV-ErbB2;Akt1^f/f^;R26Cre^ERT2^* vs. *MMTV-ErbB2;R26Cre^ER^*. P=0.0018, *MMTV-ErbB2;Akt2^f/f^;R26Cre^ERT2^* vs. *MMTV-ErbB2;R26Cre^ERT2^*. **e.** Table summarizing the incidence of lung metastasis. **f.** Circulating levels of insulin in the absence of or after tamoxifen injection. Data are presented as the means ± SEM. P=0.0005, *MMTV-ErbB2;Akt2^f/f^;R26Cre^ERT2^* vs. *MMTV-ErbB2;R26Cre^ERT2^*. **g.** Quantification of phosphorylated PRAS40, GSK3b, and Akt1(pSer473) relative to total PRAS40, GSK3b, and Akt1 in tumor extracts after tamoxifen injection into *MMTV-ErbB2;R26Cre^ERT2^* or *MMTV-ErbB2;Akt2^f/f^;R26Cre^ERT2^* mice. Data are presented as the means ±SEM. P<0.03 using an unpaired t test. **h**. Kaplan-Meier plot of tumor-free survival after tamoxifen injection into one-month-old *MMTV-NIC;Akt2^f/f^;R26Cre^ERT2^* mice. **i.** Quantification of phosphorylated Akt1 (ser473), phosphorylated pan-Akt (ser473), phosphorylated PRAS40, and phosphorylated GSK3b, relative to total Akt1, total pan-Akt, total PRAS40, and total GSK3 in tumor extracts. Data are presented as the means ±SEM. P<0.05 using an unpaired t test.

**Figure 3:**
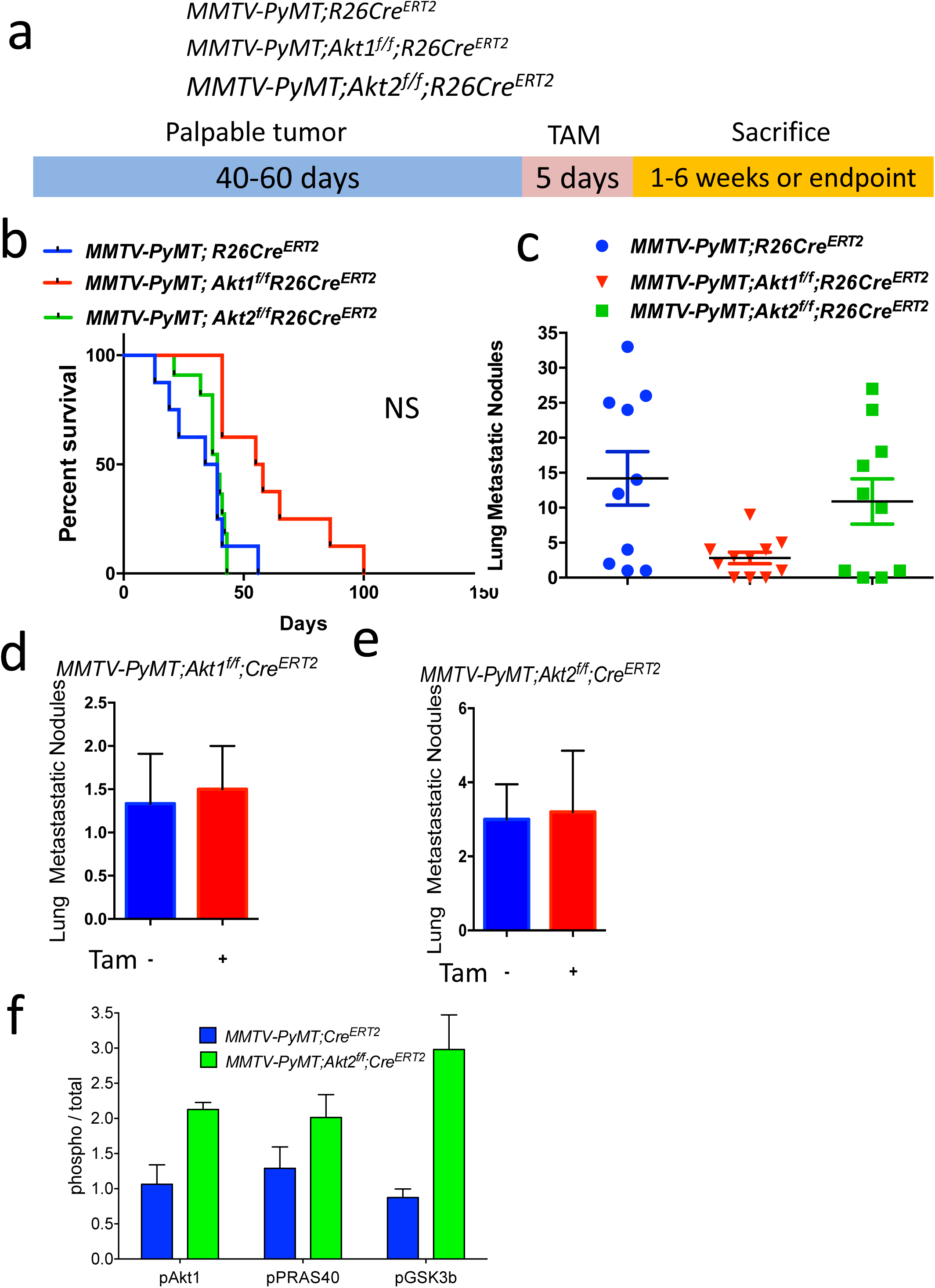
Consequences of systemic deletion of Akt1 or Akt2 after tumor onset in *MMTV-PyMT* mice. **a.** The genotypes of mice and experimental strategy. **b.** Kaplan-Meier survival curves as determined by the tumor endpoint. p=0.0092, *MMTV-PyMT;Akt1^f/f^;R26Cre^ERT2^* vs. *MMTV-PyMT;R26Cre^ERT2^*. **c.** Quantification of lung metastatic nodules. Data are presented as the means ± SEM. P=0.0066, *MMTV-PyMT;Akt1^f/f^;R26Cre^ERT2^* vs. *MMTV-PyMT;R26Cre^ERT2^* **d.** Quantification of lung metastatic nodules after orthotopic transplantation of *MMTV-PyMT;Akt1^f/f^;R26Cre^ERT2^* cells into NOG mice in the presence or absence of Akt1. Data are presented as the means ± SEM. P=0.919, using an unpaired t test. **e.** Quantification of lung metastasis nodules after the orthotopic transplantation of *MMTV-PyMT;Akt2^f/f^;R26Cre^ERT2^* cells into NOG mice in the presence or absence of Akt2. Data are presented as the means ± SEM. P=0.789, using an unpaired t test. **f.** Quantification of phosphorylated PRAS40, GSK3b, and Akt1 relative to total PRAS40, GSK3b, and Akt1 in tumor extracts after tamoxifen injection into *MMTV-PyMT;R26Cre^ERT2^*, *MMTV-PyMT;Akt1^f/f^;R26Cre^ERT2^*, and *MMTV-PyMT;Akt2^f/f^;R26Cre^ERT2^* mice. Data are presented as the means ± SEM. P<0.05, using an unpaired t test.

To analyze metastasis, the primary tumors were allowed to grow to endpoint, and subsequently the incidence of metastasis was determined. Thus, the incidence of metastasis is not a consequence of primary tumor growth. Interestingly, despite attenuating tumor development, the cell autonomous deletion of Akt1, unlike the germ line deletion of Akt1 ^5, 6^, did not inhibit the incidence of metastasis to the lung of tumor bearing mice (Fig. 1f).

## Consequences of the systemic deletion of Akt1 or Akt2 after tumor onset in *MMTV-ErbB2* mice

The cell autonomous deletion of Akt1, unlike the Akt1 germ line deletion, did not affect tumor metastasis, raising the possibility that the effect of Akt1 on metastasis is systemic. To assess the effect of the systemic deletion of different Akt isoforms after tumor formation and to emulate single isoform inhibitor drug therapy, we used *MMTV-ErbB2* mice ^8^, in which ErbB2 is overexpressed in the epithelial cells of the mammary gland. We bred *MMTV-ErbB2* mice with either *Akt1^f/f^;Rosa26(R26)Cre^ERT^*^2^ mice or *Akt2^f/f^*;*R26Cre^ERT2^* mice to generate *MMTV-ErbB2;Akt1^f/f^;R26Cre^ERT2^ and MMTV-ErbB2;Akt2^f/f^;R26Cre^ERT2^* mice (Fig. 2a). As we have shown previously, the use of *R26Cre^ERT2^* mice, in which CreERT2 was inserted in the ubiquitously expressed ROSA26 locus, enables the systemic deletion of Akt isoforms in adult mice after the injection of tamoxifen ^9^. After a latency period of 40-50 weeks, the mice developed palpable mammary tumors (approx. 0.1 cm^3^). We then performed the intraperitoneal (IP) injection of tamoxifen for 5 consecutive days to induce Akt1 or Akt2 deletion (Fig. 2a). We continuously monitored tumor growth and sacrificed the mice when the humane endpoint criteria were reached. The systemic Akt1 deletion markedly attenuated primary tumor growth (Fig. 2b, c) and extended survival (Fig. 2d). Importantly, unlike the cell autonomous Akt1 deletion in mammary gland epithelial cells, which had no effect on tumor metastasis, the systemic Akt1 deletion completely inhibited metastasis (Fig. 2e). Taken together, the results strongly suggest that the inhibition of metastasis by the systemic deletion of Akt1 is not cell autonomous.

In contrast to the cell autonomous deletion of Akt2 and the systemic deletion of Akt1, the systemic deletion of Akt2 did not inhibit tumor growth (Fig. 2b, c), but rather increased tumor growth (Fig. 2b, c), decreased survival (Fig. 2d), and markedly increased the incidence of metastasis induced by ErbB2 (Fig. 2e). We speculated that the high level of circulating insulin induced by the systemic deletion of Akt2 (Fig. 2f) hyperactivates the other Akt isoforms and thus curbs the ability of the Akt2 deletion to inhibit metastasis. Consistently, we found that the systemic deletion of Akt2 markedly elevated Akt1 phosphorylation and total Akt activity, as measured by the phosphorylation of its substrates GSK3b and PRAS40. (Fig. 2g and Supplementary Fig. 2a). Thus, the systemic deletion of Akt2 blunts the cell autonomous anti-tumorigenic activity of Akt2 (Fig. 1a) and is even pro-tumorigenic. To confirm this assertion, we crossed *MMTV-NIC;Akt2^f/f^* mice, which are resistant to tumorigenesis, with *Akt2^f/f^;R26Cre^ERT2^* mice to generate *MMTV-NIC;Akt2^f/f^;R26Cre^ERT2^* mice. Akt2 was systemically deleted in these animals at one month of age, which is more than 20 weeks before tumors were detected in *MMTV-NIC* mice (Fig. 1a). The systemic deletion of Akt2 abrogated the resistance to tumorigenesis exerted by the cell autonomous deletion of Akt2 (Fig. 2h). Importantly, the systemic deletion of Akt2 elevated Akt1 and possibly Akt3 activities and enabled the expression of ErbB2 in the absence of Akt2 in the mammary glands of *MMTV-NIC;Akt2^f/f^;R26Cre^ERT2^* mice (Fig. 2i, and supp. Fig. 2b), which is not expressed after cell autonomous deletion of Akt2 (Fig. 1c). These results provide strong additional support for our assertion that ErbB2 expression in the mammary gland is not tolerated when Akt activity is below a certain threshold level.

To determine the cell autonomous effect of Akt1 and Akt2 after tumor formation, the tumor cells derived from late stage tumors in *MMTV-ErbB2;Akt1^f/f^;Cre^ERT2^* or *MMTV-ErbB2;Akt2^f/f^;R26Cre^ERT2^* mice were orthotopically transplanted into nonobese diabetic (NOD)/Shi-scidIL-2Rγ^null^ (NOG) mice. When the tumors were palpable, the mice were exposed to tamoxifen for 5 consecutive days to delete Akt1 or Akt2. The deletion of Akt1 markedly attenuated tumor growth (Supp. Fig. 3a), whereas the deletion of Akt2 attenuated tumor growth to a much lesser extent (Supp. Fig. 3b). The relative effect of Akt1 versus Akt2 on tumor growth is directly correlated with their relative individual expression in late stage tumors in which Akt1 expression is induced and Akt2 expression declines (Fig. 1e). Indeed, when compared to the Akt2 deletion, the deletion of Akt1 in *MMTV-ErbB2* orthotopic tumors markedly decreased the total level of Akt, indicating that Akt1 is the predominant isoform at late stages of tumor development (Supp. Fig. 3c).

## Consequences of the systemic deletion of Akt1 or Akt2 after tumor onset in *MMTV-PyMT* mice

Polyoma virus middle T-antigen expression in the mammary gland of mouse mammary tumor virus-polyoma middle tumor antigen (*MMTV-PyMT)* mice induces several signaling pathways that are altered in human breast cancer, including the SRC and PI3K pathways. Specifically, the MMTV-PyMT mouse model will result in the development of multifocal mammary adenocarcinomas with a high incidence of metastatic lesions to the lymph nodes and lungs ^10^. Therefore, we employed this mouse model to study the effect of systemic Akt1 or Akt2 deletion on the incidence of metastasis. We generated *MMTV-PyMT;R26Cre^ERT2^, MMTV-PyMT; Akt1^f/f^;R26Cre^ERT2^* and *MMTV-PyMT;Akt2^f/f^;R26Cre^ERT2^* mice. After tumor onset, the systemic deletion of Akt1 or Akt2 was induced by tamoxifen injection. These mice were followed and subsequently sacrificed at the endpoint to assess metastasis (Fig. 3a). The Akt1 systemic deletion significantly increased tumor free survival, whereas the Akt2 systemic deletion did not (Fig. 3b). The systemic Akt1 deletion markedly reduced the number of metastatic nodules in the lungs but the systemic Akt2 deletion did not (Fig. 3c). To further establish that the effect of Akt1 on metastasis is not cell autonomous, we orthotopically implanted cells derived from tumors in *MMTV-PyMT; Akt1^f/f^;R26Cre^ERT2^* and *MMTV-PyMT;Akt2^f/f^;R26Cre^ERT2^* mice into NOG mice. After the tumors became palpable, the mice were treated with tamoxifen for 5 consecutive days. When tumors reached the endpoint, the mice were analyzed for metastases in the lungs. As shown in Fig. 3d and 3e, the cell autonomous deletion of neither Akt1 nor Akt2 changed the number of metastatic nodules, further supporting the systemic effect of Akt1 on metastasis. These results are consistent with the results obtained in *MMTV-ErbB2* mice and further confirmed the discrepancy between the cell autonomous and systemic effects of Akt isoforms on mammary gland tumorigenesis. Indeed, similar to the results found in *MMTV-ErbB2;Akt2^f/f^;R26Cre^ERT2^* mice, the systemic deletion of Akt2 in *MMTV-PyMT;Akt2^f/f^;R26Cre^ERT2^* mice hyperactivated Akt1 and total Akt activity in the tumors (Fig. 3f). The discrepancy between the inducible cell autonomous and non-cell autonomous effects of Akt1 and Akt2 deletions on tumor growth and metastasis again suggested that the inhibition of Akt1 might be superior to the inhibition of Pan-PI3K or Pan-Akt as a tumor therapy.

## Systemic Akt1 deletion inhibits metastasis by impairing neutrophil mobilization

To further delineate the systemic effects of Akt1 and Akt2 on mammary gland tumorigenesis and metastasis, we adopted Drop-seq technology for single cell RNA sequencing (scRNA-seq) as previously described ^11^. Using this approach, we sequenced 7,791 cells over 5 biological replicates from the primary tumors of *MMTV-PyMT* mice and identified 17 clusters (Fig. 4a, Supplementary Fig. 5a, Supplementary Table 1). Surprisingly, we identified nine distinct clusters of cells within the primary tumors based on the expression of *PyMT* (Supplementary Fig. 5b, and Fig. 4), indicating that the tumors were very heterogeneous. Previous studies showed that keratin 14 (Krt14) is a marker for disseminating early metastatic cells, and its deletion suppresses metastasis^12^. Interestingly, within the nine distinct clusters of cells, we found one cluster (cluster 13) that expressed high levels of Krt14 (Fig. 4b). Cluster 13 also expressed other epithelial markers, such as *Krt5*, *Krt7*, *Krt8*, *Krt17*, and *Krt18*. However, cluster 13 also expressed a relatively high level of Vimentin (*Vim*), an EMT marker. Previous studies have shown that the high expression of Krt14 is also associated with the high expression of genes that regulate metastasis, such as Tenascin C (Tnc), Adam metallopeptidase (Adamts1), Caveolin 1 (Cav1), Jagged1 (Jag1), and Proepiregulin (Ereg) ^12^. Indeed, cluster 13 expressed high levels of *Tnc* and *Adamts1*, as well as *Cav1*, *Jag1*, and *Ereg* (Fig. 4b and Supplementary Table 1). Notably, cluster 11, which expressed high levels of vimentin, also expressed high levels of *Ereg* and *Adamts1* and relatively high levels of *Cav1* and *Jag1* (Fig. 4b). However, high *Krt14* expression, unlike in cluster 13, was not found in cluster 11. Paradoxically, cluster 13 expresses the highest level of E-cadherin (*cdh1*), whereas cluster 11 expresses the lowest level of *cdh1* (Supplementary Table 1). However, this is consistent with a recent report showing that E-cadherin is required for the survival of disseminating metastatic cells in this mouse model ^13^.

**Figure 4.**
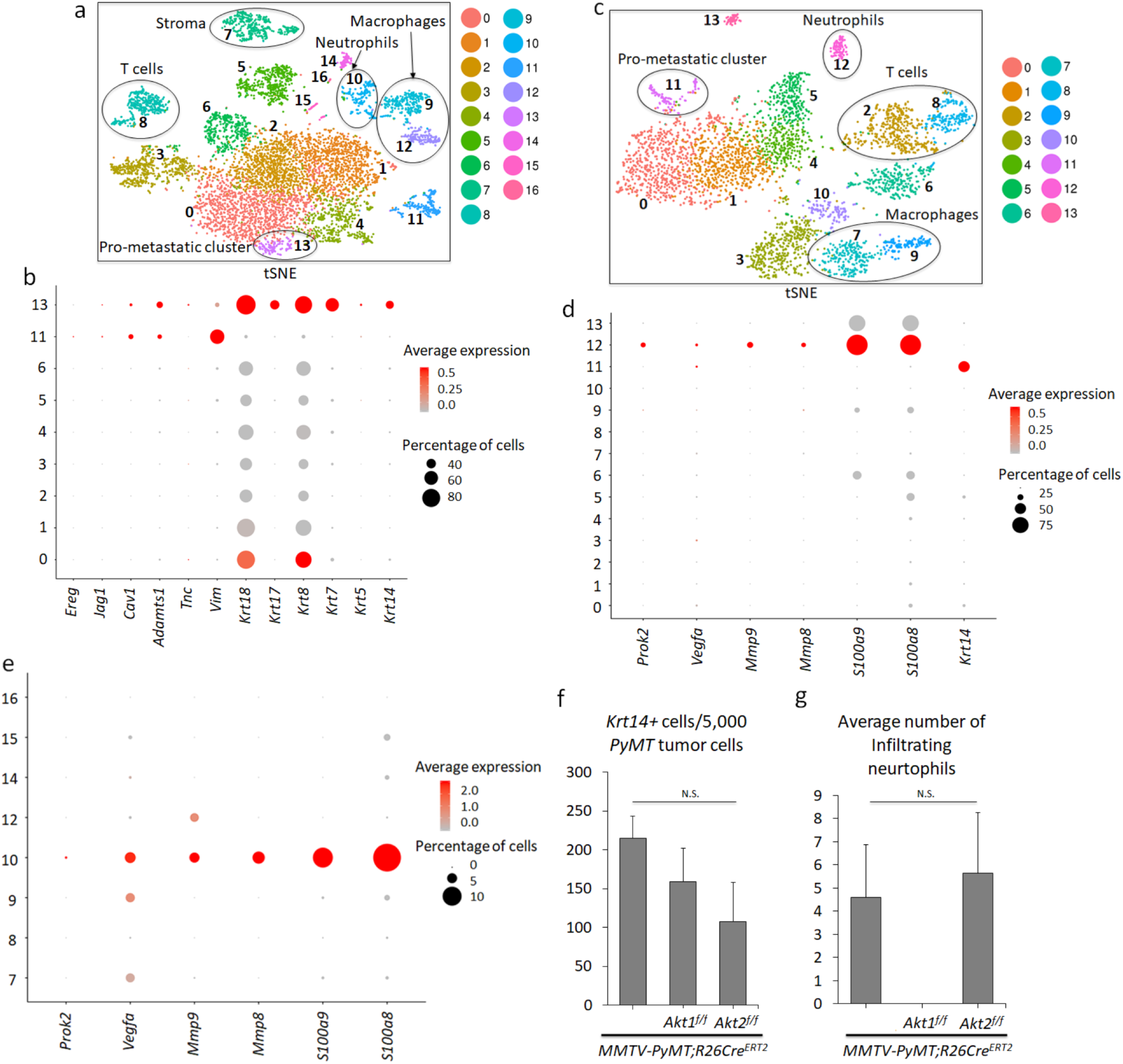
Analysis of primary and metastatic tumors by scRNA-seq. **a.** t-Distributed Stochastic Neighbor Embedding (t-SNE) plot of primary mammary gland tumors in *MMTV-PyMT* mice. Each cluster is characterized by a unique gene expression signature. A total of 7,791 primary breast tumor cells (N = 5) were used. Clusters 0, 1, 2, 3, 4, 5, 6, 11, and 13 are tumor cells expressing *PyMT,* whereas the remaining clusters are non-tumor cells. The clusters are color-coded. **b.** Dot plot showing expression of metastatic markers *Ereg, Jag1, Cav1, Adamts1, Tnc, Vim, as well as Krt18, Krt17, Krt8, Krt7, Krt5,* and *Krt14* across the *PyMT* primary tumor clusters. **c.** tSNE plot of 3,979 metastatic tumor cells in the lung (N = 3). Clusters 0, 1, 4, 5 and 11 are *PyMT*-positive. **d.** Dot plot depicting expression of *Prok2, Vegfa, Mmp9, Mmp8, S100a9, S100a8,* and the prometastatic marker *Krt14* across the clusters in the lung metastatic scRNA-seq analysis. **e.** Dot plot in the primary non-tumor clusters indicate that cluster 10 has a predominant gene expression profile of neutrophil markers *Prok2, Vegfa, Mmp9, Mmp8, S100a9, S100a8*. **f.** Graph showing the average number of pro-metastatic *Krt14*-positive cells for every 5,000 *PyMT*-positive tumor cells based on the scRNA-seq results. N(*MMTV-PyMT*) = 5, N(*MMTV-PyMT; Akt1^f/f^*) = 3, N(*MMTV-PyMT; Akt2^f/f^*) = 3. N. S = not significant (p>0.05). One-way analysis of variance (ANOVA) was used to calculate significance. **g.** Graph showing the average number of infiltrating neutrophils in each genotype. scRNA-seq reveals the absence of neutrophil cells in primary systemic *Akt1* knockout tumors across all 3 biological replicates. N(*MMTV-PyMT*) = 5, N(*MMTV-PyMT; Akt1^f/f^*) = 3, N(*MMTV-PyMT; Akt2^f/f^*) = 3. N.S = not significant (p>0.05). One-way ANOVA was used to calculate significance. Error bars represent standard error.

Thereafter, we sequenced 3,979 single cell transcriptomes over 3 biological replicates from the macroscopic metastatic lesions in the lungs of *MMTV-PyMT* mice (Fig. 4c). The scRNA-seq results revealed five clusters (0, 1, 4, 5, and 11) that were derived from the primary tumors as determined by the high level of *PyMT* expression (Supplementary Fig. 5c, Supplementary Table 2). Among the five clusters, only one, cluster 11, expressed high levels of *Krt14* (Fig. 4d). After combining and re-clustering the primary and metastatic scRNA-seq results, we found that cluster 19 (Supplementary Fig. 6, and supp. Table 4) consisted of cells from both the metastatic lung cluster 11 (Fig. 4c) and primary mammary tumor cluster 13 (Fig. 4a). This finding indicates that the last two clusters are similar and share a gene expression profile and traces a metastatic cluster found in the lung to a cluster in the primary tumors.

Micrometastases express high levels of Krt14, which are diminished in macrometastases ^12^. Therefore, we analyzed the lungs of tumor-bearing mice that do not display macroscopic metastases. Consistently, we found only one cluster derived from the primary tumors, and this cluster expressed high levels of *Krt14* and *PyMT* (Supplementary Fig. 5d, e, Supplementary Table 3). Thus, among the distinct clusters in the primary tumors, we classified the high *Krt14*-expressing cluster as a pro-metastatic cluster.

Within the primary tumor, we also found nontumor cells that did not express *PyMT*, which include stromal cells, T cells, macrophages, and neutrophils (Fig. 4a). Importantly, a population of neutrophils was also found within the metastatic tumors in the lungs (Fig. 4c). Neutrophils play a pro-metastatic role in breast cancer ^14, 15^, and a high neutrophil to lymphocyte ratio (NLR) is associated with worse overall survival and disease-free survival ^16^. Consistent with the reported pro-metastatic role of neutrophils that provide a niche for the metastatic cells in the lungs, we found that neutrophils within lung metastases and primary tumors express, in addition to high Ly6g and Cxrc2, which are known neutrophils markers, relatively high levels of *S100a8*, *S100a9*, *MMP8*, *MMP9*, *Bv8*/*Prok2* and vascular endothelial growth factor (*Vegfa*), which promote invasion and migration (Fig. 4d, e, and Supplementary Fig. 5e).

After the systemic deletion of Akt1 or Akt2, analysis of the tumors using Drop-seq (3,194 cells over 3 replicates and 4,647 cells over 3 replicates, respectively) revealed that primary tumors had similar cell clusters as those in control mice, including a prometastatic cluster expressing high Krt14, which is co-segregated in control wild type (WT), Akt1-/-, and Akt2-/- primary tumors (Supplementary Fig. 7, and supp. Table 5). Furthermore, the percentage of high Krt14-expressing cells in primary tumors after the systemic deletion of Akt1 or Akt2 was not significantly different from that in control primary tumors (Fig. 4c). These results suggest that the systemic Akt1 deletion did not change the relative presentation of the pro-metastatic cluster within the primary tumors.

While the presentation of a high Krt14-expressing cell cluster was similar in WT control mice and in mice after the systemic deletion of Akt1 or Akt2 (Fig. 4f, and supp. Table 5), the presentation of neutrophils was completely diminished after Akt1 systemic deletion (Fig. 4g, and supp. Fig. 7). These results raised the possibility that systemic Akt1 deletion inhibits metastasis by impairing neutrophil mobilization to the lung. To further assess this possibility, we examined whether systemic Akt1 deletion could affect pro-metastatic neutrophils in the lungs. The percentage of neutrophils in the lungs of non-tumor-bearing mice, as measured by anti-Ly6G staining, was not significantly different in control mice and in mice after either Akt1 or Akt2 systemic deletion (Fig. 5a). However, in tumor-bearing mice, the percentage of neutrophils in the lungs of control mice or in mice after Akt2 systemic deletion was markedly increased, whereas Akt1 systemic deletion did not increase the percentage of neutrophils in the lungs (Fig. 5b).

**Figure 5.**
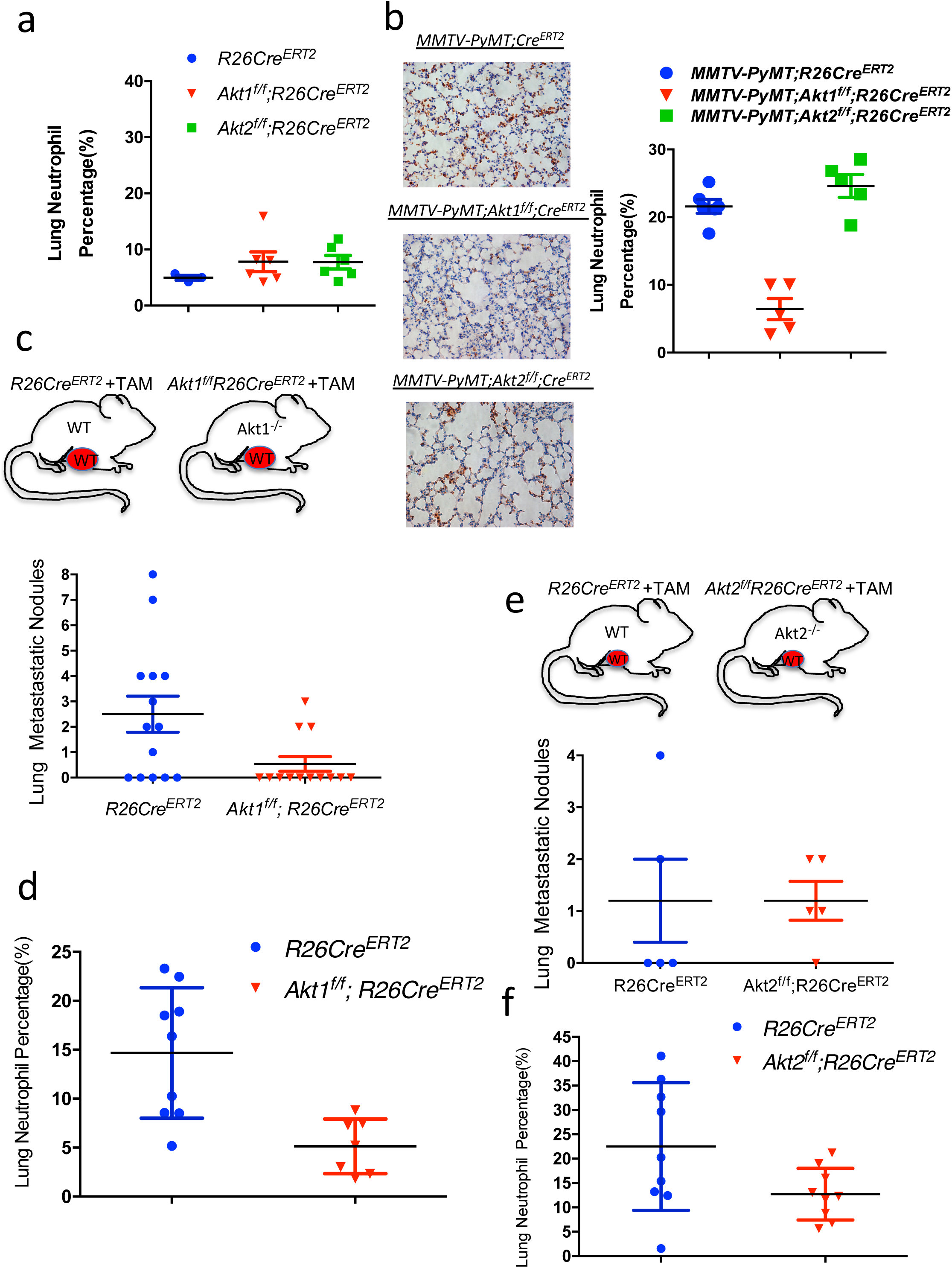
The effect of the systemic Akt1 or Akt2 deletion on neutrophil accumulation in the lungs of tumor-bearing mice. **a.** Percentage of neutrophils in the lungs of nontumor-bearing mice after tamoxifen injection to systemically delete Akt1 or Akt2. The percentage of neutrophils was calculated by counting the number of Ly6G-positive cells relative to total hematoxylin and eosin-stained cells in lung tissue at endpoint sections as described in the Methods section. Data are presented as the means +/- SEM. P= 0.3, using unpaired t test. **b.** Left: Quantification of lung neutrophils in control *MMTV-PyMT* mice and after systemic deletion of Akt1 or Akt2. Right: Representative lung section images stained with anti-Ly6G. Quantification was performed after the primary tumors reached the endpoint. Data are presented as the means ± SEM. P=0.0066, *MMTV-PyMT;Akt1^f/f^;R26Cre^ERT2^* vs. *MMTV-PyMT;R26Cre^ERT2^* and p=0.518, *MMTV-PyMT;Akt2^f/f^;R26Cre^ERT2^* vs. *MMTV-PyMT;R26Cre^ERT2^* using an unpaired t test. Quantification of metastatic nodules (**c**) and neutrophils (**d**) in the lungs of *R26Cre^ERT2^* and *Akt1^f/f^;R26Cre^ERT2^* mice orthotopically transplanted with MMTV-PyMT tumor cells and injected with tamoxifen at palpation. Quantification was performed when the primary tumors reached the endpoint. Data are presented as the means ±SEM. P=0.019 for metastatic nodules, and P=0.0034 for neutrophils using an unpaired t test. Quantification of metastatic nodules (**e**) and neutrophils (**f**) in the lungs of *R26Cre^ERT2^* and *Akt2^f/f^;R26Cre^ERT2^* mice orthotopically transplanted with MMTV-PyMT tumor cells and injected with tamoxifen at palpation. Quantification was performed when the primary tumors reached the endpoint. Data are presented as the means ±SEM. P>0.05 for metastatic nodules and neutrophils using an unpaired t test.

If systemic Akt1 deletion inhibits metastasis by a systemic effect that impairs neutrophil mobilization to the lungs, then this deletion would also inhibit the metastasis of WT tumors. We therefore orthotopically implanted tumor cells derived from *MMTV-PyMT* mice into either *R26Cre^ERT2^*, *Akt1^f/f^;R26Cre^ERT2^*, or *Akt2^f/f^;R26Cre^ERT2^* mice. When the tumors became palpable, the mice were injected with tamoxifen to systemically delete Akt1 or Akt2. When the tumors reached the endpoint, the mice were analyzed for metastasis. The systemic deletion of Akt1 markedly decreased the metastasis of WT tumors (Fig. 5c), which is directly correlated with the decrease in tumor-associated neutrophils in the lungs (Fig. 5d). However, when compared to WT control mice, the systemic deletion of Akt2 did not decrease the metastasis of WT tumors (Fig. 5e) and did not significantly affect the percentage of neutrophils in the lungs (Fig. 5f).

If the Akt1 deficiency in neutrophils determines decreased metastatic potential, then it is expected that the specific deletion of Akt1 in neutrophils could decrease metastasis. Therefore, we used *MRP8-Cre* mice ^17^ to delete Akt1 specifically in neutrophils. We orthotopically implanted E0771 mouse breast cancer cells into control *MRP8-Cre*, and *MRP8-Cre;Akt1^f/f^* mice. As shown in Fig. 6a, lung metastasis was diminished in *MRP8-Cre;Akt1^f/f^* mice compared to that of *MRP8-Cre* mice. Consistently, neutrophils were accumulated in the lungs of *MRP8-Cre* mice but markedly reduced in *MRP8-Cre;Akt1^f/f^* mice (Fig. 6b). Thus, these results provide direct evidence that the systemic effect of the Akt1 deletion on metastasis is due to the effect on pro-metastatic or tumor-associated neutrophils.

**Figure 6.**
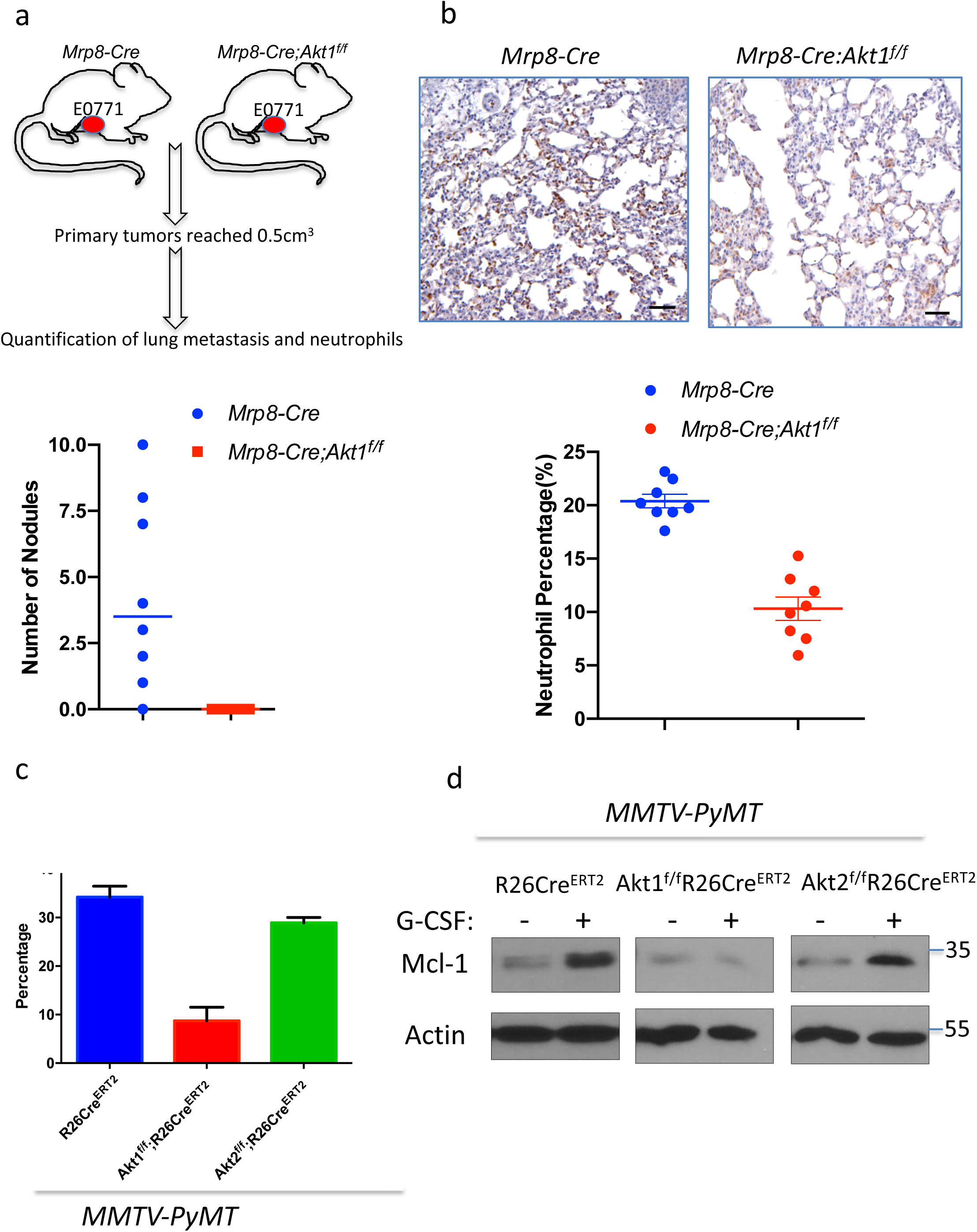
Consequences of Akt isoform deletion in the neutrophils of tumor-bearing mice. **a.** The effect of Akt1 deletion on metastasis in *MRP8-Cre* mice after orthotopic transplantation of E0771 cells. Upper panel: Schematic of experimental design. Bottom panel: Quantification of metastasis. **b.** The effect of Akt1 deletion on neutrophils in the lungs of *MRP8-Cre* tumor-bearing mice (n=8). Upper panels show representative lung section images stained with anti-Ly6G. Data are presented as the means ±SEM. P<0.0001, for metastatic nodules and neutrophils using an unpaired t test. **c.** The effect of G-CSF on the survival of neutrophils isolated from control, and systemically deleted Akt1 and Akt2 tumor-bearing mice. **d.** The effect of G-CSF on the level of MCL1 in neutrophils isolated from control and systemically deleted Akt1 and Akt2 tumor-bearing mice.

**Figure 7.**
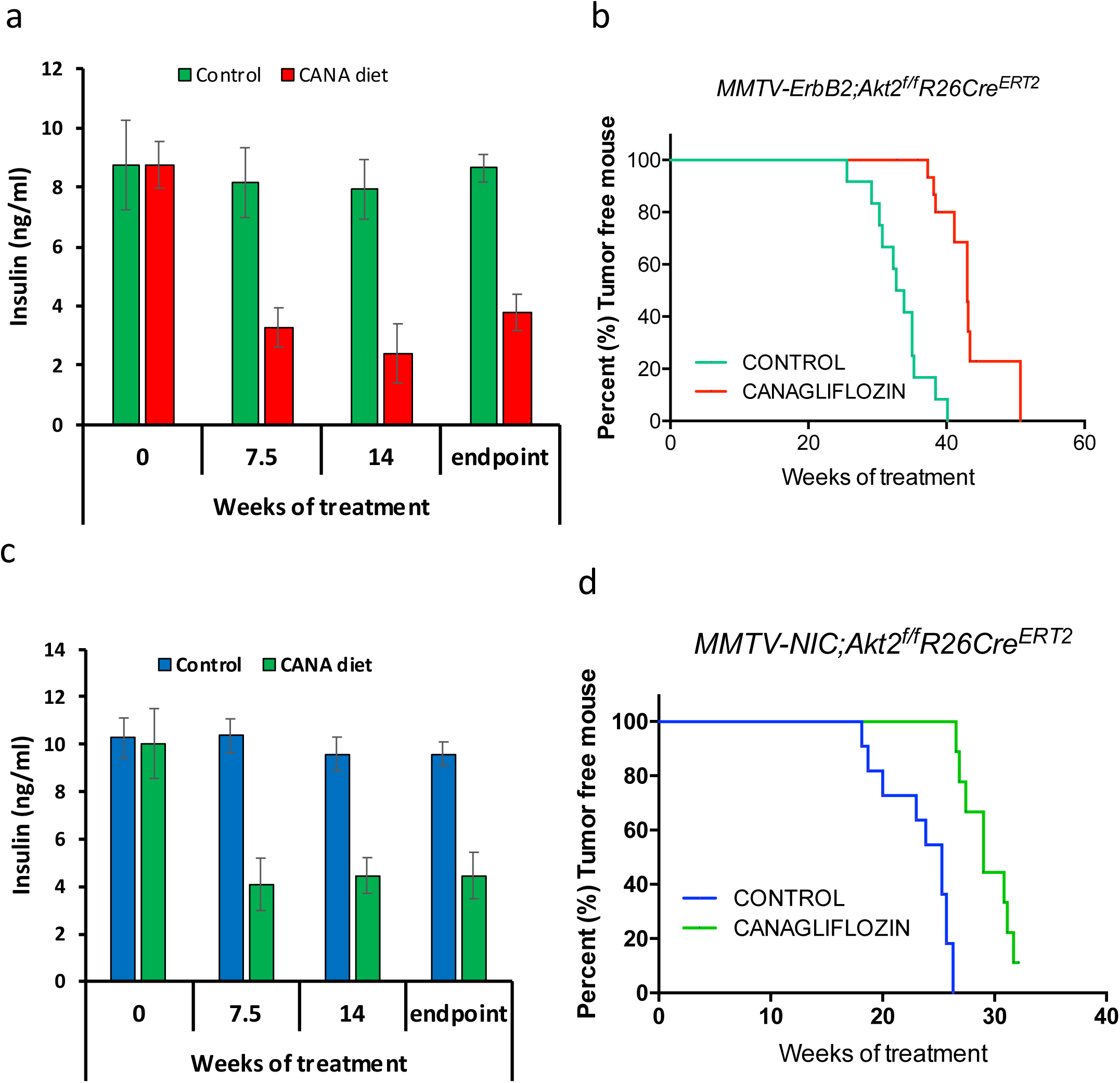
Reducing circulating levels of insulin by SGLT2 inhibitor attenuates the mammary gland tumorigenesis after systemic Akt2 deletion. *MMTV-ErbB2;Akt2^f/f^;R26Cre^ERT2^* or *MMTV-ErbB2;Akt2^f/f^;R26Cre^ERT2^* mice were injected with tamoxifen at one month of age and were subjected to either control chow diet or diet that includes canagliflozin (CANA). **a,c.** Insulin level after chow diet or CANA diet. **b,d.** Tumor onset after chow diet or CANA diet.

To understand the mechanism by which Akt1 affects tumor-associated neutrophils, we isolated neutrophils from tumor-bearing mice and exposed these cells to granulocyte colony stimulating factor (G-CSF) in vitro. G-CSF was shown to promote the survival of neutrophils and is required for the mobilization of tumor-associated neutrophils ^14^. As shown in Fig. 6c, G-CSF increased the survival of neutrophils isolated from either control or Akt2-deficient tumor-bearing mice but not the survival of neutrophils isolated from Akt1-deficient tumor-bearing mice. These results suggest that, at least in part, Akt1 deficiency decreases the survival of tumor-associated neutrophils. In mouse models of mammary gland tumors, the inhibition of translation initiation factor eIF4E decreases metastasis to the lungs by decreasing the mobilization of neutrophils to these organs ^18^. This finding was attributed to the increase in neutrophil cell survival by eIF4E after exposure to G-CSF through the elevation of BCL2 and MCL1 ^18^. Since Akt, which is upstream of mTORC1 and eIF4E ^19^, could also affect the level of MCL1 through the inhibition of GSK3 and the increase in its protein stability ^20^, we examined the level of MCL1 after exposure to G-CSF. We found that MCL1 protein levels are induced by G-CSF in neutrophils derived from either control or Akt2-deficient tumor-bearing mice but not in neutrophils derived from Akt1-deficient tumor-bearing mice (Fig. 6d). Thus, Akt1 is required downstream of G-CSF to promote elevated MCL1 protein. These results are consistent with other studies showing that MCL1 is particularly important for the survival of neutrophils and is induced by G-CSF ^21^. One possible reason why Akt1 and not Akt2 affects neutrophils in tumor-bearing mice is that Akt1 is the major expressed isoform in neutrophils of tumor bearing mice (supp. Fig. 8). Taken together, these results showed that the systemic effect of Akt1 deficiency on the inhibition of metastasis is due to the effect of Akt1 on the survival and mobilization of the neutrophils that promote metastasis.

## Reducing insulin level after systemic Akt2 deletion inhibits mammary tumor development

Our results suggested that the systemic deletion of Akt2 curbs the inhibition of mammary gland tumorigenesis because of the high circulating levels of insulin that hyperactivates the other Akt isoforms and other pro-oncogenic signaling pathways (Fig. 2 and Fig. 3). To further assess this possibility, we treated mice with the diabetic drug, metformin, to decrease insulin levels after systemic Akt2 deletion. However, metformin was not able to reduce insulin levels in these mice. We therefore fed the mice with a diet that includes the sodium-glucose co-transporter (SGLT2), inhibitor, the anti-diabetic drug, canagliflozin. SGLT2 is responsible for reabsorption of glucose in the kidney, and therefore limits the excretion of glucose through the urine ^22^. Inhibition of SGLT2 by its selective inhibitor, canagliflozin, was shown to inhibit hyperglycemia and hyperinsulinemia without adverse consequences ^22^. As shown in Fig. 2d and 2h, the systemic deletion of Akt2 in *MMTV-ErbB2* and *MMTV;NIC* mice accelerated and exacerbated tumorigenesis, which was attributed to the high level of insulin. To affirm this possibility *MMTV-ErbB2;Akt2^f/f^;R26Cre^ERT2^* or *MMTV-NIC;Akt2^f/f^;R26Cre^ERT2^* mice were fed, after systemic Akt2 deletion, with a diet containing canagliflozin, with an average approximate dose of 40mg/kg BW/day. As shown in Fig. 7a,c canagliflozin substantially reduced the elevated insulin level observed after Akt2 systemic deletion. Consequently, the development of mammary gland tumorigenesis was markedly reduced when mice were treated with canagliflozin after systemic Akt2 deletion (Fig. 7b,d). Thus, systemic deletion of Akt2 does not inhibit and may even accelerate mammary gland tumorigenesis by elevating blood insulin levels. Reducing insulin levels after systemic Akt2 deletion inhibits the acceleration of mammary gland tumorigenesis by systemic Akt2 deletion.

## Discussion

The results described here underscore the importance of employing systemic deletion as a genetic proof of concept for cancer therapy, as cell autonomous deletion could otherwise be misleading. This notion was exemplified by the effect of the cell autonomous versus systemic deletion of Akt1 or Akt2 on breast cancer development and metastasis. Surprisingly, the cell autonomous deletion of Akt1 at tumor onset and after tumor formation did not inhibit metastasis, whereas systemic deletion of Akt1 markedly inhibited metastasis. We showed that the predominant mechanism by which systemic Akt1 deletion inhibits metastasis is by inhibiting the survival and mobilization of pro-metastatic neutrophils. These neutrophils express relatively high levels of prokineticin 2 (PROK2) and VEGEFa, which promote angiogenesis, as well as the metalloproteases MMP8 and MMP9, which promote invasion. These effects would allow the extravasation and mobilization of neutrophils but could also promote cancer cell angiogenesis and invasion. Neutrophils also physically interact with cancer cells ^23^, and thus can increase the extravasation of disseminating cancer cells by expressing and secreting MMP8 and MMP9. More recently, it was shown that neutrophils actually escort disseminating cancer cells to the metastatic site ^24^; thus, it is possible that these cells also promote cancer cell extravasation. In addition, neutrophils form neutrophil extracellular traps (NETs) that stimulate migration and invasion and trap natural killer cells ^25^. The pro-metastatic role of neutrophils was also recognized in human cancer patients, and a high NLR is associated with a poor prognosis ^14^. The neutrophils derived from tumor-bearing mice or from cancer patients are distinct from normal neutrophils, as tumor-associated neutrophils lack immunosuppressive activity and have a higher migration capacity ^26^. Cancer cells promote the survival and mobilization of neutrophils by secreting G-CSF ^14, 27^. We showed that at least in vitro Akt1, and not Akt2, is required for the G-CSF-induced survival of neutrophils derived from tumor-bearing mice. The major mechanism by which G-CSF promotes the survival of neutrophils is by inducing the expression of the anti-apoptotic protein MCL1 ^21^. We showed that Akt1 deficiency prohibits the induction of MCL1 expression by G-CSF. We cannot completely exclude, however, other potential mechanisms by which the Akt1 deficiency impairs the survival and mobilization of tumor-associated neutrophils. Notably, we found that Akt1 is the major expressed Akt isoform in neutrophils of tumor bearing mice and this could explain why systemic deletion of Akt1and not Akt2 markedly affect the pro-metastatic neutrophils. Previous studies have shown the differential roles of Akt isoforms in neutrophil function. In contrast to our findings with respect to neutrophil function in tumor-bearing mice, the neutrophils derived from nontumor-bearing mice are more dependent on Akt2 than on Akt1 in response to stimulation by N-Formylmethionyl-leucyl-phenylalanine (fMLP) or phorbol myristate acetate (PMA) ^28^. This finding could be explained by the change in the phenotype of these neutrophils in response to tumor formation and by the different stimuli used. Finally, It was speculated that in tumor bearing hosts there is a pressure to release neutrophils from the bone marrow prematurely and that these immature neutrophils can be converted to pro-tumorigenic pro-metastatic neutrophils ^29^. We identified cluster 18 in supp. Fig. 6a and 6b, or cluster 21 in supp. Fig. 7a,c as neutrophils that are also present in the lung of tumor bearing mice but are missing after systemic deletion of Akt1 (Fig. 4g, and supp. Fig. 7c). These neutrophils were identified as the pro-metastatic neutrophils and express the highest RNA level of Ly6g and Cxcr2, which are known neutrophil markers. However, we also identified cluster 14 in supp. Fig. 7a, c, as a potential neutrophil progenitor population. This cluster had the second highest RNA expression of Cxcr2 with no expression of Ly6g. It is tempting to speculate that Akt1 might be required for the differentiation of the progenitor neutrophils population at the primary tumor site, but more work beyond the scope of this manuscript is required to assess this possibility. Nevertheless, there is no difference in the percentage of cells in cluster 14 between control mice and Akt1 or Akt2 deleted mice (supp. Fig. 7c).

Our studies also show that the expression of ErbB2 in the absence of Akt2 only in mammary gland cells cannot be tolerated and these cells are likely eliminated. We hypothesized that ErbB2 expression cannot be tolerated in mammary gland cells in which Akt activity is below a certain threshold level. Akt2 is expressed at the highest level whereas Akt1 is expressed at the lowest level in early stages of mammary gland tumorigenesis. Thus, it is possible that total Akt activity is more reduced at this stage in the absence of Akt2 than in the absence of Akt1. However, systemic Akt2 deletion, which increases insulin levels and hyperactivates Akt1 and possibly Akt3, overcomes the intolerance of ErbB2 expression in the absence of Akt2 in the mammary gland. In late stage tumor growth, Akt1 expression is elevated, and Akt2 expression declines; thus, the growth of tumor cells derived from *MMTV–ErbB2* mice is impaired to a much higher extent by Akt1 deletion than by Akt2 deletion. However, it remains to be determined how the cell autonomous deletion of Akt1, which is expressed at the highest level in late stages of tumor development, is tolerated in the presence of high ErbB2 expression. One possibility is that at late stages, the cells have already acquired additional lesions that enable the expression of ErbB2, despite low Akt activity.

The systemic deletion of Akt2 markedly increased tumor growth and metastasis in *MMTV-ErbB2* mice and did not inhibit tumor growth and metastasis in *MMTV-PyMT* mice. This effect was attributed to the high circulating levels of insulin as a consequence of systemic Akt2 deletion. Indeed, reducing insulin levels after the systemic deletion of Akt2 inhibits tumor onset. Recently it was shown that reducing insulin level after treatment with pan-PI3K inhibitors, which elevate insulin level, increased their efficacy ^30^. Our results showed that downstream of PI3K inhibition, Akt2 inhibition is responsible for the elevated insulin.

Taken together, these results provide strong support for the use of systemic deletion as a proof of concept for cancer therapy. These results together with our previous results ^9, 31^ provide support for using specific Akt1 inhibitors and avoiding Akt2 or pan-Akt inhibitors for cancer therapy. Furthermore, the effect of Akt1 systemic deletion on pro-metastatic neutrophils, but not on other functions of neutrophils, indicate that Akt1 specific inhibitor would be sufficient to selectively inhibit the pro-metastatic effect of neutrophils.

## Methods

### Mouse strains and mouse work

The *MMTV-NIC* mice and *MMTV-Erbb2* mice were gifts from W.J. Muller (McGill University). The *MMTV-PyMT* and *LSL-Luc* mice were purchased from Jackson Laboratory. The FVB/N WT mice were purchased from Charles River Laboratories. The *R26Cre^ERT2^* knockin mice (strain 01XAB) Akt1^f/f^, and Akt2^f/f^, *Akt1^f/f^;R26RCre^ERT2^* and *Akt2^f/f^;R26RCre^ERT2^* mice have been previously described ^9^. *MMTV-PyMT;Akt1^f/f^;R26RCre^ERT2^* and *MMTV-PyMT;Akt2^f/f^;R26RCre^ERT2^* mice were generated by crossing *Akt1^f/f^;R26RCre^ERT2^* or *Akt2^f/f^;R26RCre^ERT2^* mice with *MMTV-PyMT* mice. *MMTV-Erbb2;Akt1^f/f^;R26RCre^ERT2^* and *MMTV-Erbb2;Akt2^f/f^;R26RCre^ERT2^* were generated by crossing *Akt1^f/f^;R26RCre^ERT2^* or *Akt2^f/f^;R26RCre^ERT2^* mice with *MMTV-Erbb2* mice. *MMTV-NIC*;*Akt1^f/f^* and *MMTV-NIC;Akt2^f/f^* mice were generated by crossing *Akt1^f/f^* or *Akt2^f/f^* with *MMTV-NIC* mice. *MMTV-NIC;Akt2^f/f^;R26RCre^ERT2^* mice were generated by crossing *MMTV-NIC;Akt2^f/f^* mice with *Akt2^f/f^;R26RCre^ERT2^* mice. *MMTV-NIC;LSL-Luc* mice were generated by crossing MMTV-NIC mice with LSL-Luc mice. *MMTV-NIC;Akt1^f/f^;LSL-Luc* mice and *MMTV-NIC;Akt2^f/f^;LSL-Luc* were generated by crossing of *MMTV-NIC;LSL-Luc* mice with *Akt1^f/f^* and *Akt2^f/f^* mice respectively. All mouse models were produced in an FVB background. Mice from other backgrounds were backcrossed to FVB/N mice for at least 10 generations. Cre recombinase activation was performed by the IP injection of 0.1 ml of 20 mg/ml of tamoxifen for 5 consecutive days. For IP injection, tamoxifen was dissolved in corn oil at a final concentration of 20 mg/ml via shaking at 37°C for 30 minutes as previously described ^9^. PCR, qPCR and western blot analysis of multiple tissues, including tumor tissues, were used to confirm the deletion of Akt isoforms. C57Bl6 *MRP8-Cre-ires-GFP* mice were purchased from Jackson Laboratory and were crossed with *Akt1^f/f^* mice in C57Bl6 background to generate *MRP8-Cre;Akt1^f/f^* mice. NOG mice were purchased from Jackson Laboratory. For the Canagliflozin diet experiments, the mice were injected with tamoxifen at 6 weeks of age and maintained on chow diet (Teklad #7012). On day 15, blood samples were collected from the tail vein using a heparinized micro-capillary tube to measure basal fed plasma glucose and insulin. Mice were then randomly divided into two groups. Canagliflozin (CANA) (MedChem Express, Monmouth Junction, NJ, USA) was administrated as a food additive at a concentration of 0.03% (w/w) into Teklad #7012 chow diet (Research Diets, Inc., New Brunswick, NJ, USA). Each group received control or CANA diet with ad libitum access to food until the day they are sacrificed. The average daily dose of canagliflozin (calculated from average daily food intake for mice and actual body weight) was approximately 40mg/kg BW/day, a dose effective in many studies ^32–34^. The body weight of each mouse was measured every week. Fed plasma glucose and insulin levels were measured every 4 weeks. Starting at 6-month of age, mice were palpated each week for tumor appearance and age of appearance was recorded. All animal experiments were approved by the Institutional Animal Care and Use Committee of the University of Illinois at Chicago (UIC), as required by the United States Animal Welfare Act and the policy of NIH.

### Primary tumor cell isolation

Mouse tumor tissues were dissected, washed in PBS supplemented with 5% P/S (penicillin/streptomycin), cut into small pieces (diameter∼3 mm), and then digested in 1% collagenase IV in DMEM at 37°C for 30 minutes with shaking. The supernatant was discarded after centrifugation at 300 g for 5 minutes. The pellet was washed several times with PBS and passed through a 75 μm cell strainer to collect small cell clumps. The clumps were then transferred to plates containing DMEM supplemented with 10% FBS and 1% P/S in an incubator at 37°C with 5% CO_2_.

### Tumor cell transplantation

Freshly isolated tumor cells (less than 3 days) were harvested and resuspended in PBS and Matrigel (Corning) in a 1:1 ratio with a final concentration of 0.1∼1 x 10^6^/100 μl. One hundred microliters of cells were transplanted to the fourth mammary gland fat pad of each side to recipient mice. Mice were monitored/palpated daily for 1 week to confirm successful tumor transplantation. After the tumor diameter reached 4 mM, the tumor size was measured every week until a certain time point or a humane endpoint. Tumor sizes were measured with a caliper and calculated by length*height*width*0.5. For transplantation E0771 mouse breast cancer cells, 1x10^5^ cells were orthotopically transplanted into the mammary glands of *MRP8-Cre* and *MRP8-Cre;Akt1^f/f^* mice as described above. When primary tumors reached 0.5cm^3^ the mice were analyzed for lung metastasis and neutrophils infiltration.

### Bone marrow isolation

After muscle removal, the femurs and tibias were collected and rinsed with 70% ethanol followed by ice-cold PBS wash. The epiphyses were cut off. Then, a 10-cc syringe filled with RPMI medium supplemented with 10% FBS and 2 mM EDTA was used to flush the marrow cells from both ends with a 25-gauge needle into a 50 ml Falcon tube with a 40 μm cell strainer. After centrifugation at 1000 rpm for 5 min, the pellet was resuspended in 3 mL cold ammonium chloride potassium (ACK) lysis buffer for 1 min, centrifuged and washed with PBS and resuspended in the desired volume.

### Neutrophil isolation

Bone marrow-derived neutrophils were isolated from mice using the following protocol adapted from Swamydas *et al*. ^6^. Bone marrow cells were obtained from mice by flushing the contents of the tibia and femur with complete RPMI medium supplemented with 2 mM EDTA using a 25-gauge needle. The cell suspension was run through a 100 µM cell strainer. The cells were pelleted at 430xg for 7 minutes at 4°C. Red blood cells were lysed by washing the cells with 20 mL of 0.2% NaCl for 20 seconds followed by the addition of 20 mL of 1.6% NaCl. After washing with PBS, the bone marrow cells were resuspended in ice-cold PBS and layered on the top of a Histopaque 1119/1077 (Sigma) gradient in a 15 mL Falcon tube. After centrifugation at 2000 rpm at room temperature for 30 minutes (without the break), the neutrophils were isolated by collecting the cells at the interface of the two Histopaque layers. The neutrophils were then washed twice with complete RPMI medium. Neutrophil viability and purity were assessed via trypan blue staining and flow cytometric analysis of Ly6G surface expression by using a PE-Ly6G antibody (Biolegend).

### Cell culture

Freshly isolated mouse breast tumor cells were cultured in high glucose DMEM supplemented with 10% FBS and 1% P/S. Freshly isolated neutrophils were cultured in RPMI 1640 supplemented with 10% FBS and 1% P/S.

### Neutrophil cell death assay

Freshly isolated neutrophils were cultured with or without G-CSF at a final concentration of 100 ng/ml for 24 hours. The neutrophils were then stained with Hoechst stain and propidium iodide (PI) for 30 minutes in an incubator followed by plate scanning with a Celigo Imaging Cytometer. The dead cell/live cell ratio was determined by the red cell number (PI stained)/blue cell number (Hoechst stained) ratio.

### Measurement of glucose, insulin, and IL-6 levels

Glucose levels were measured with a glucometer test strip (Precision Xtra; Abbott Laboratories). Insulin levels were measured by Milliplex immunoassay (Millipore) according to the manufacturer’s instructions.

### Tissue staining and immunohistochemistry

Breast tumor tissues were freshly collected and directly fixed in 10% formalin (Fisher Chemical). The lungs were inflated with PBS via the trachea and then removed from the ribcage, washed in PBS and dissected to lobules, before fixing in 10% formalin. After fixation for 24-48 hours (depending on tissue size) in formalin, the tissues were processed and embedded in paraffin blocks. The sections (5 μm) were stained with hematoxylin and eosin (H&E). For immunohistochemistry, antigen retrieval was performed by incubating the sections in 0.01 M sodium citrate (pH 6.0) at 95°C for 20 minutes, followed by cooling down to room temperature. The sections were treated with 0.3% H_2_O_2_ for 5 minutes to quench endogenous hydrogen peroxide. After blocking with normal serum, the sections were incubated with primary antibody at 4°C overnight. After incubation with the appropriate secondary antibodies from Vectastain ImmPress Kits (Vector Labs), a 3, 3 -diaminobenzidine (DAB) kit (Vector Labs) was applied to visualize the signal. The sections were then lightly counterstained with hematoxylin. To determine the lung metastatic incidence metastatic nodules were counted under the microscope after H&E staining. Infiltration of neutrophils to the lungs was quantified histology staining for Ly6G antibody. Pictures of 5 random fields were taken from each lung section. Quantification was performed via ImageJ. Lung neutrophil percentage = Ly6G positive cell number / total cell number in the lung.

### Protein analysis by immunoblotting

The cells were harvested and washed with cold PBS and lysed in lysis buffer [20 mM HEPES, 1% Triton X-100 (TX-100), 150 mM NaCl, 1 mM EGTA, 1 mM EDTA] containing phosphatase inhibitors (10 mM sodium pyrophosphate, 20 mM β-glycerol-glycerophosphate, 100 mM sodium fluoride, 5 mM indoleacetic acid, 20 nM oleic acid) and a Pierce^TM^ protease inhibitor mini tablet inhibitor and a Pierce^TM^ phosphatase inhibitor mini tablet inhibitor (1 tablet per 10 mL lysis buffer). After sonication on ice, the solubilized proteins were collected by centrifugation, and the protein concentration was measured via a Bio-Rad protein assay. An equal amount of protein was aliquoted into Laemmli buffer, boiled for 5 minutes and processed by standard western blot procedures. The membranes were blocked with 5% skim milk in TBS with 0.1% Tween-20 for 1 hour at room temperature and incubated with specific primary antibodies at 4°C overnight. Enhanced chemiluminescence (ECL) western blot substrate was used for film development, and ImageJ was used for data quantification. The tissue samples collected from the animal were snap frozen in a dry ice-ethanol mixture and preserved at -80°C until needed for the experiment. The tissue samples for western blotting were homogenized in ice-cold lysis buffer and processed as stated above.

### Single cell RNA-seq

Mouse tumor tissues were dissected, washed in PBS, cut into small pieces and then digested in 1% collagenase IV in DMEM at 37 < for 45 minutes with shaking. Cell debris and red blood cells were removed with centrifuging and ACK lysis buffer. The cell pellets were washed with PBS and passed through a 40 μm cell strainer to form a single cell suspension and counted with trypan blue to determine cell number and viability. A sequencing library was constructed according to the Drop-Seq protocol^11, 35^. Briefly, approximately 1.5-2.0x10^5^ cells were resuspended in 1 mL 0.1% BSA PBS and loaded onto a microfluidic chip to isolate single cell droplets. The collected droplets were broken to release barcoded beads and resuspended in reverse transcriptase mix. After reverse transcription, the beads were treated with exonuclease I to remove unhybridized DNA, followed by PCR amplification. The amplified products were sent to the UIC Research Resources Center (RRC) to check the cDNA quality and quantity with TapeStation and Qubit. The libraries were constructed using the Nextera XT kit and then sequenced with the Illumina Nextseq 500 at RRC. The reading depth was approximately 40,000 per cell. Read 1 was 20 base pairs (bp) to determine the cell barcode and unique molecular identifier (UMI); read 2 was 63 bp to determine the cDNA region.

### Bioinformatics data processing and analysis

Raw sequence data were filtered, trimmed and then aligned to the mouse genome (mm10). Uniquely mapped reads were grouped and counted to generate digital expression matrices and subjected to Seurat package (version v2.2.1) using R (version 3.3.2) to perform a single cell analysis^11, 35^.

### Materials

|  |  |  |
| --- | --- | --- |
| Xylenes (Histological), Fisher Chemical | Fisher Scientific | Cat. # X3S-4 |
| Hematoxylin Solution, Harris Modified | Sigma-Aldrich | Cat. # HHS32-1L |
| Eosin Y solution, alcoholic | Sigma-Aldrich | Cat. # HT110116 |
| Recombinant Mouse G-CSF Protein | R&D Systems | Cat. # 414-CS-025 |
| Pierce™ ECL Western Blotting Substrate | Thermo Fisher Scientific | Cat. # 32106 |
| Pierce™ Protease Inhibitor Mini Tablets | Thermo Fisher Scientific | Cat. # A32953 |
| Pierce™ Phosphatase Inhibitor Mini Tablets | Thermo Fisher Scientific | Cat. # A32957 |
| HyClone™ Dulbecco's High Glucose Modified Eagles Medium | Fisher Scientific | Cat. #SH30022.01 |
| HyClone™ RPMI 1640 Media | Fisher Scientific | Cat. #SH30027.01 |
| HyClone™ Phosphate Buffered Saline (PBS) | Fisher Scientific | Cat. #SH30256.01 |
| Propidium iodide | Sigma-Aldrich | Cat. # P4170-25MG |
| Hoechst 33342, trihydrochloride, trihydrate | Thermo Fisher Scientific | Cat. # H3569 |
| Histopaque®-1077 | Sigma-Aldrich | Cat. # 10771-100ML |
| Histopaque®-1119 | Sigma-Aldrich | Cat. # 11191-100ML |
| Gibco™ ACK Lysing Buffer | Fisher Scientific | Cat. # A1049201 |
| Corning Matrigel Matrix | Corning Life Sciences | Cat. # 354230 |
| Collagenase, Type 4 | Worthington | Cat. #LS004188 |
| Tamoxifen | Sigma-Aldrich | Cat. #T5648-5G |
| Corn Oil | Sigma-Aldrich | Cat. #C8267-500ML |
| MK-2206 | Selleck Chem | Cat. #S1078 |

### Commercial kit Assays

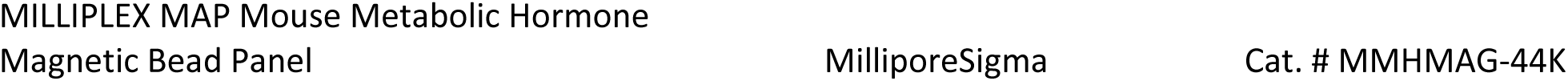

### Antibodies

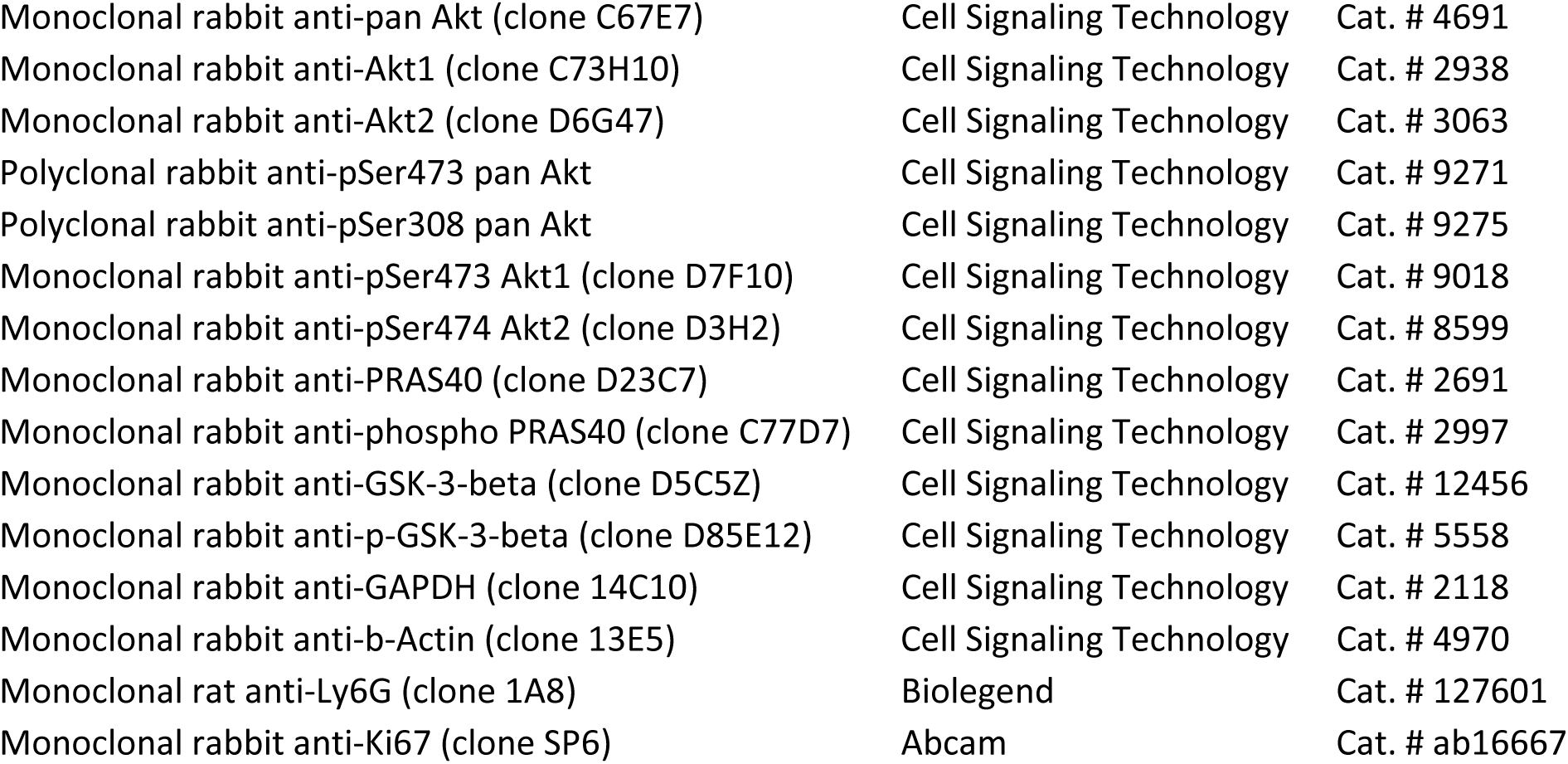

## Acknowledgments

N.H. acknowledges the support from NIH grants R01AG016927, R01CA090764, and R01 CA206167, the VA merit award BX000733, and the VA research career scientist award IK6BX004602. X.C. acknowledges the support from the center for clinical and translational science. C.B acknowledges the support from F30CA228191. W.P. acknowledges the support from T32HL007829. M.M.A acknowledges the support from the center for clinical and translational science. M.V.F. acknowledges the support from NIH grants R01GM093827 and R35GM131707. We would like to thank Ling Jin for maintaining and genotyping the mice. N.H. would like to thank Louis Philipson (University of Chicago) for the suggestion to use SGLT2 inhibitors.

## Author Contributions

N.H. and X.C. conceived the study. N.H., X.C., G.R., V.N. C.B. and W.P. designed the experiments. X.C. generated the mouse models and performed most of the experiments. M.M.A and X.C. conceived the Drop-seq experiments. M.M.A performed and analyzed the Drop-seq experiment and supervised by M.V.P. and A.B.M.M.K.I. G.R. designed and performed the experiments in Fig. 1d and experiments with Mrp8-Cre mice. V.N. designed and performed the experiments in Fig. 2h, 2i, and Fig. 7. W.P. analyzed neutrophils from tumor bearing mice. N.H., X.C. and M.M.A wrote the manuscript.

## Supplementary figure legends

**Supplementary Figure 1:**
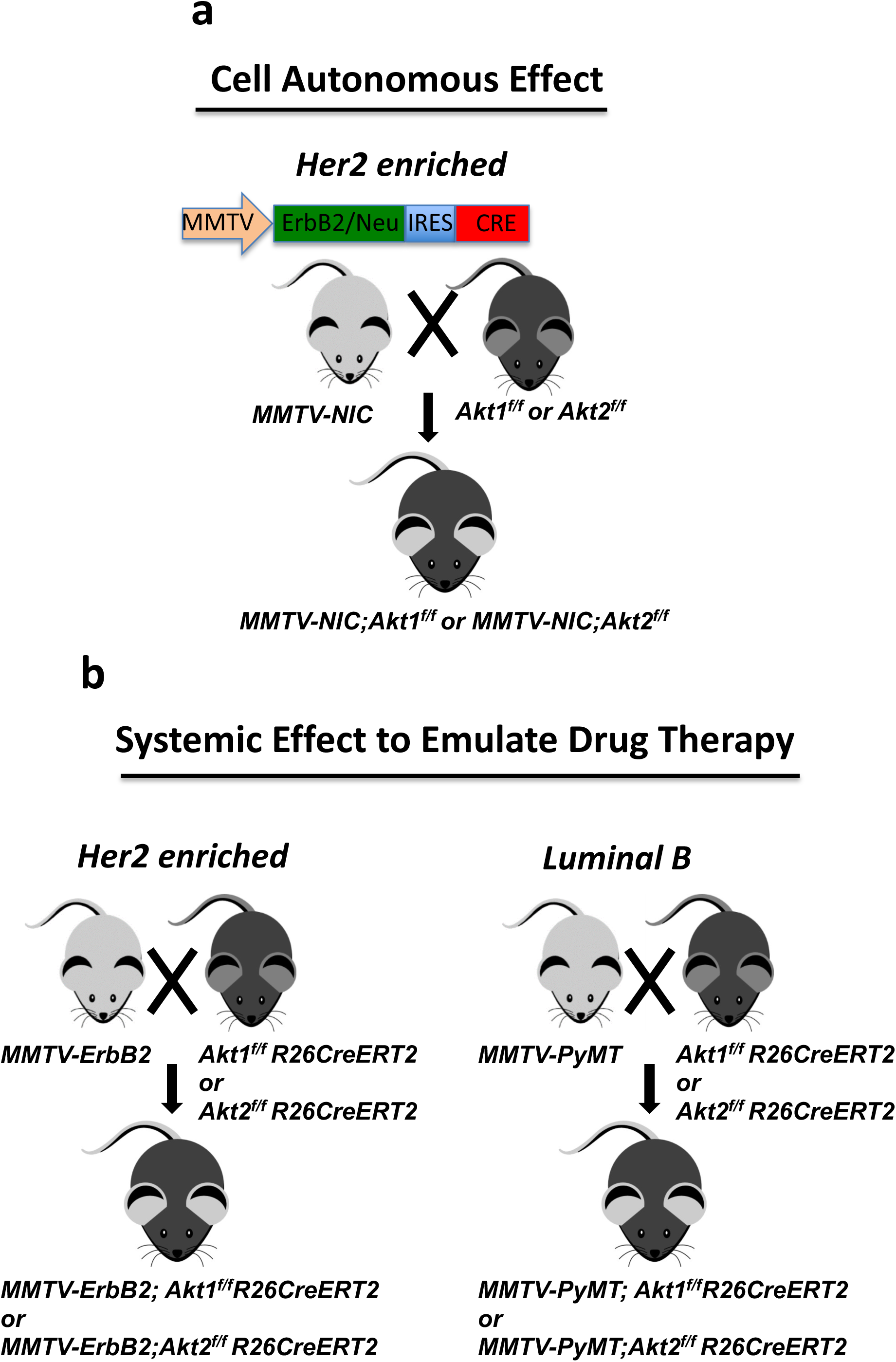
Breeding schemes. **a.** Breeding to generate a high Her2 mouse model with mammary gland-specific deletion of Akt1 or Akt2 to determine the cell autonomous effect. **b.** Breeding to determine the systemic effect of Akt1 or Akt2 deletion in high Her2 (left panel) and luminal B (right panel) mouse models.

**Supplementary Figure 2a:**
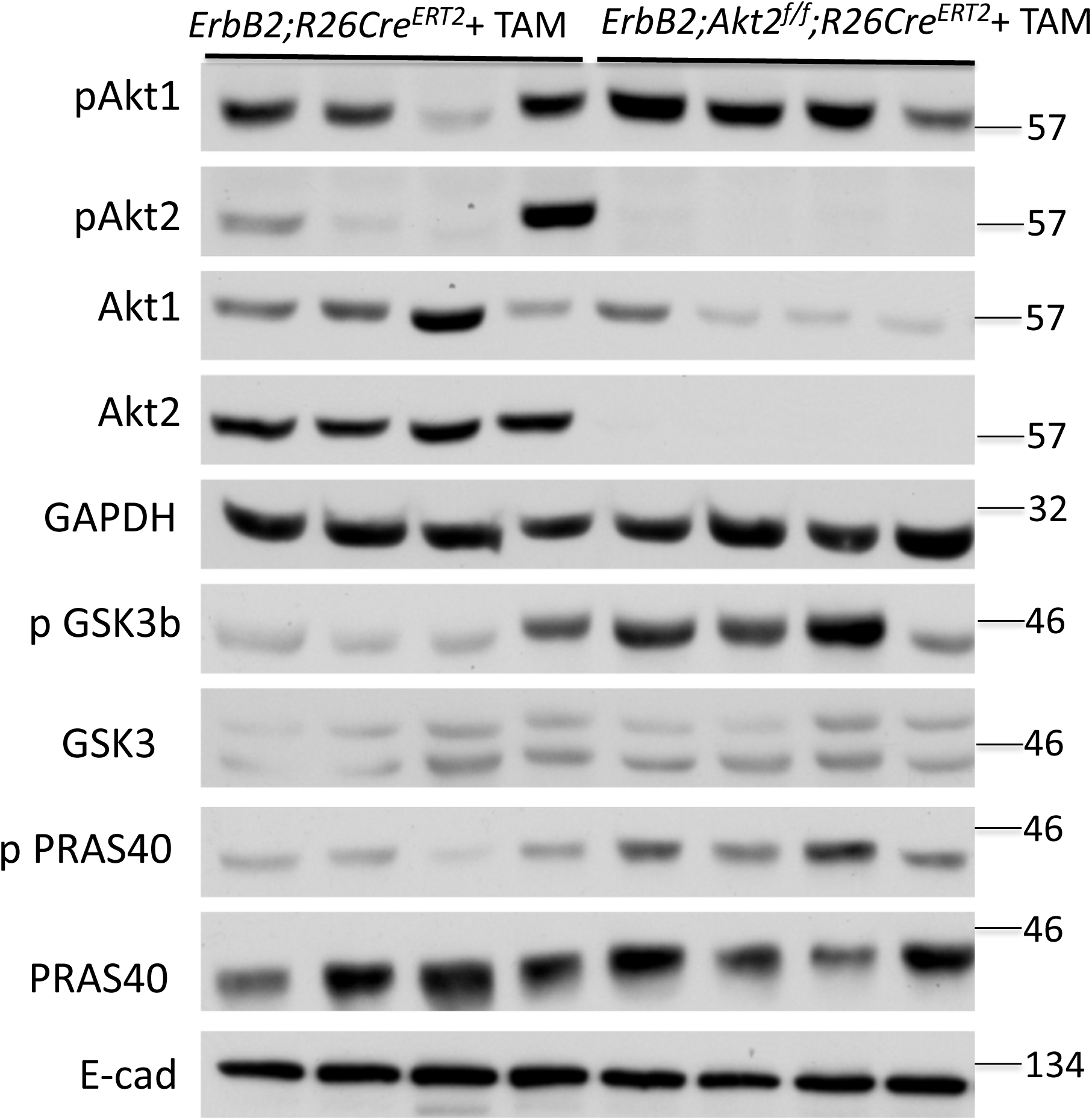
Representative immunoblot showing the phosphorylation of Akt1 (pSer473), GSK3b, and PRAS40 in mammary gland tumors derived from control *MMTV-ErbB2;R26Cre^ERT2^* or *MMTV-ErbB2;Akt2^f/f^;R26Cre^ERT2^* mice after the systemic deletion of Akt2. Extracts from individual tumors in four different mice were used. Immunoblots were used for quantifications in Fig. 2g.

**Supplementary Figure 2b:**
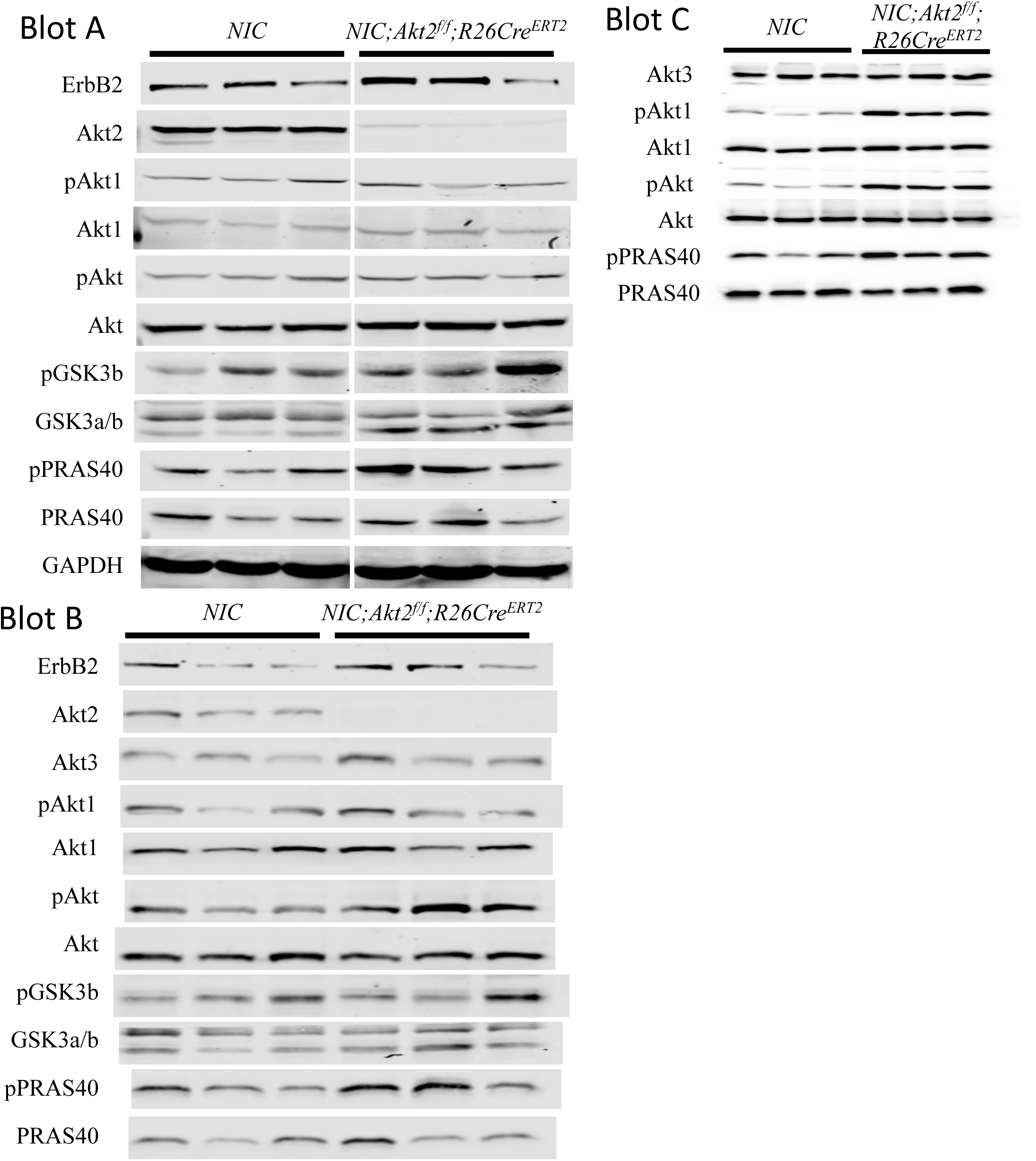
Representative immunoblots showing expression of Akt1, Akt2, Akt3, and total Akt, GSK3b, PRAS40, phosphorylation of Akt1 (pSer473), phosphorylation of pan-Akt (pSer473), phosphorylation of GSK3b, and PRAS40 in mammary gland tumors derived from control *MMTV-NIC or MMTV-NIC;Akt2^f/f^;R26Cre^ERT2^* mice after the systemic deletion of Akt2. Extracts from individual tumors in nine different mice were used. Immunoblots were used for quantifications in Fig. 2i.

**Supplementary Figure 3:**
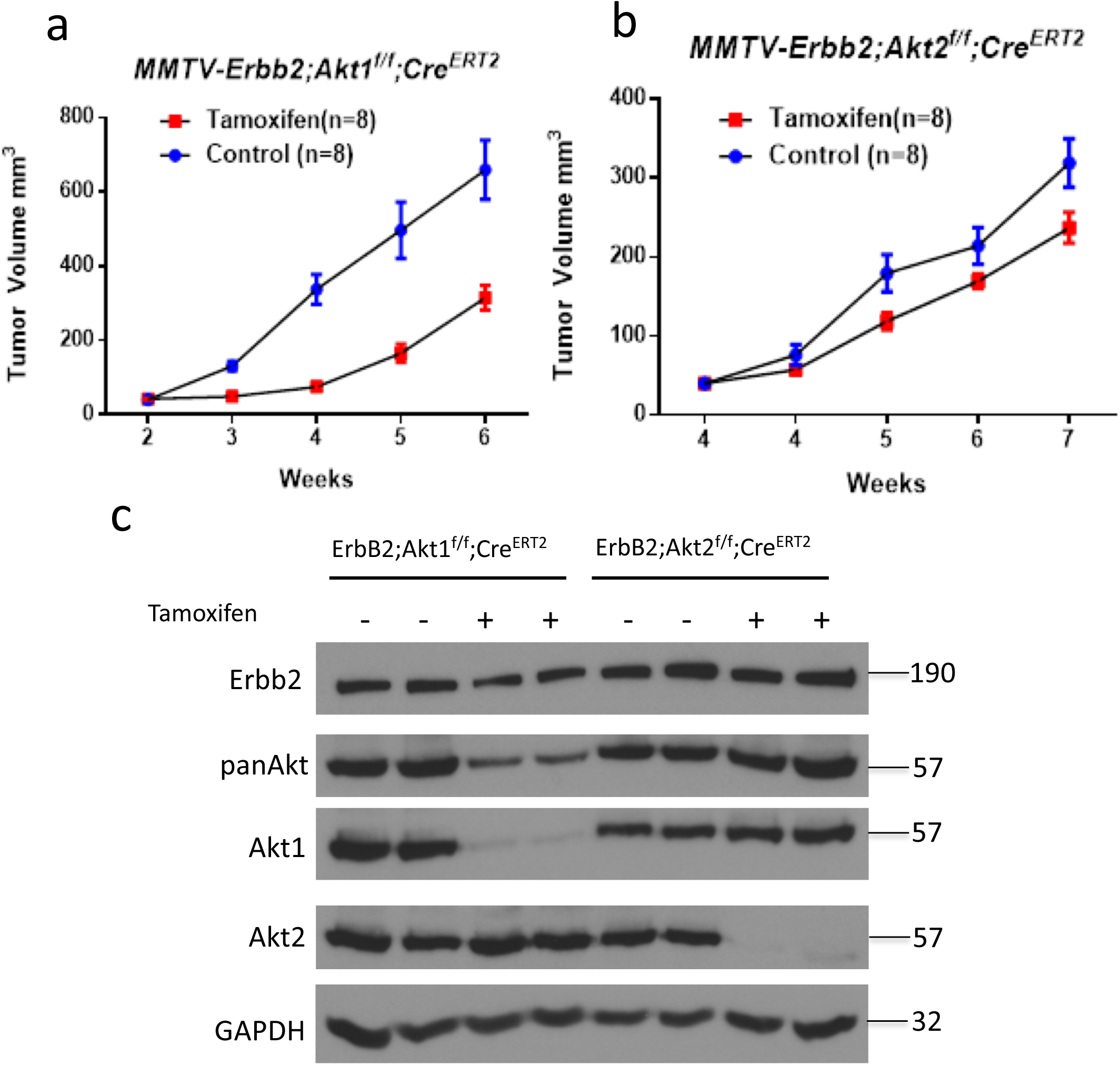
Cell autonomous deletion of Akt1 or Akt2 in orthotopically transplanted tumors in NOG mice. **a.** Tumor growth curve of mammary tumor cells derived from *MMTV-ErbB2;Akt1^f/f^;R26RCre^ERT2^* mice and orthotopically implanted into the mammary fat pad of NOG mice. After palpation, the mice were either treated or not treated with tamoxifen to delete Akt1. Cell autonomous Akt1 deletion significantly impaired tumor growth (n=8, p<0.001). **b.** Tumor growth curve of mammary tumor cells derived from *MMTV-ErbB2;Akt2^f/f^;R26RCre^ERT2^* mice and orthotopically implanted into the mammary fat pad of NOG mice. After palpation, the mice were either treated or not treated with tamoxifen to delete Akt2. Cell autonomous Akt2 deletion significantly impaired tumor growth (n=8, p<0.001). **c.** Representative immunoblot showing the level of total Akt expression in orthotopic tumors after the deletion of Akt1 or Akt2.

**Supplementary Figure 4:**
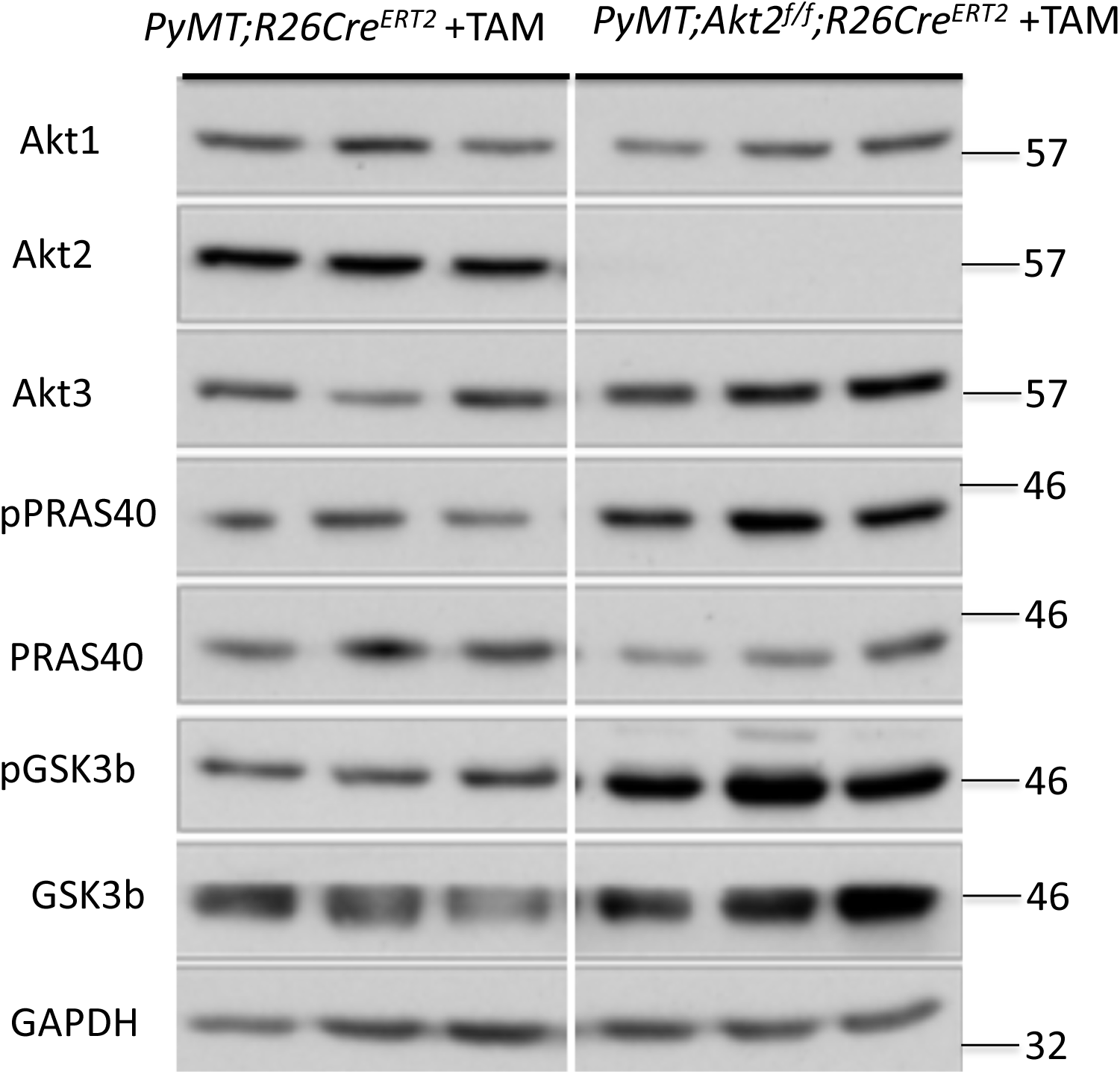
Representative immunoblot showing the phosphorylation of GSK3b and PRAS40 in mammary gland tumors derived from control *MMTV-PyMT;R26Cre^ERT2^* mice or after the systemic deletion of Akt2. Extracts from individual tumors in three different mice were used.

**Supplementary Figure 5:**
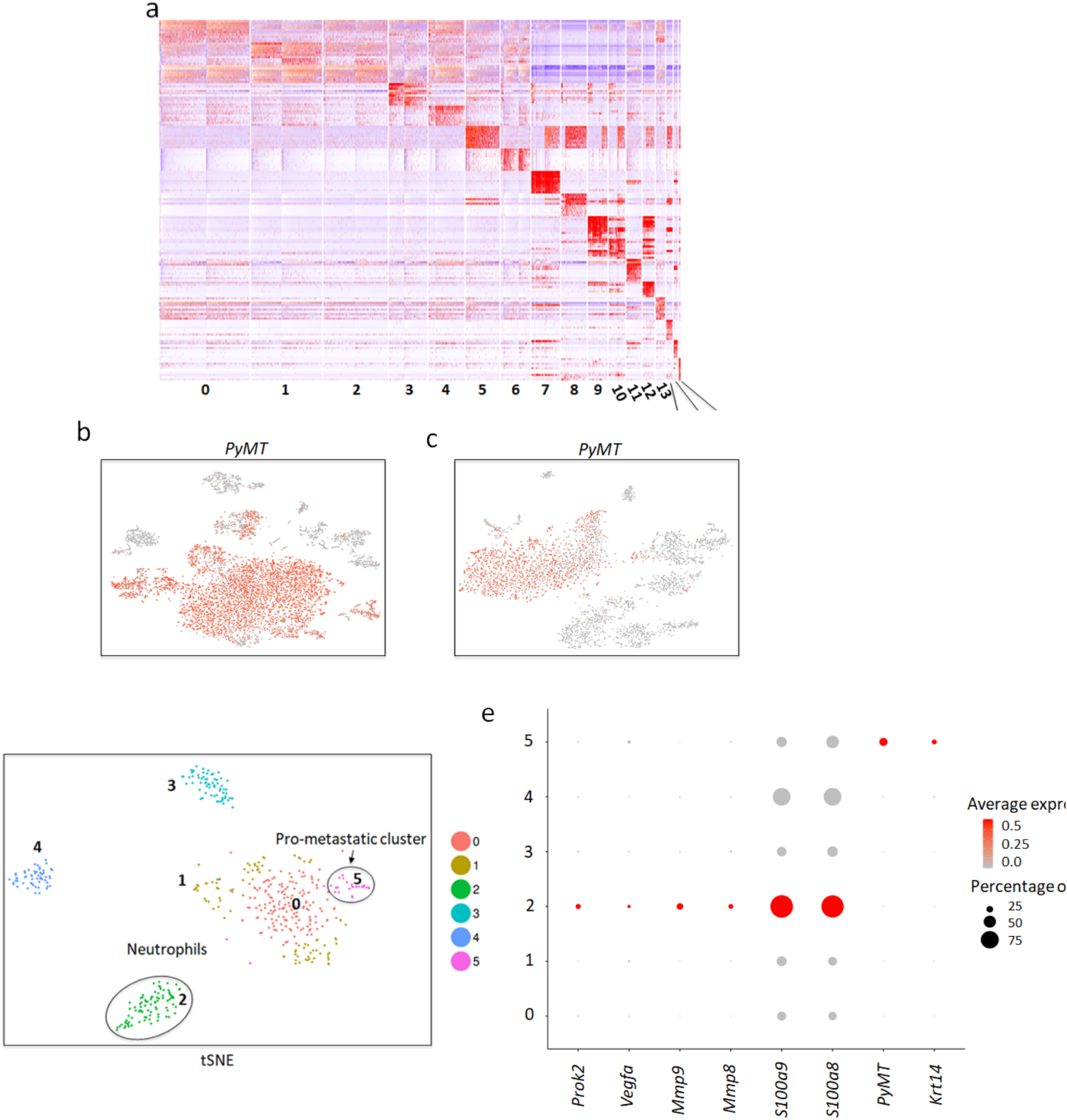
**a.** Heatmap showing distinct markers in each cluster using scRNA-seq on the primary *MMTV-PyMT* breast tumor tissue. **b.** Feature plot showing the cells expressing *PyMT* in the primary breast tumor analysis. **c.** Feature plot showing the cells expressing *PyMT* in the metastatic lung tumor analysis. **d.** Analysis of lung micrometastatic lesions revealing prometastatic cluster 5. **e.** Dot plot showing the expression of *PyMT* and *Krt14* in cluster 5 only in the micrometastasis analysis.

**Supplementary Figure 6:**
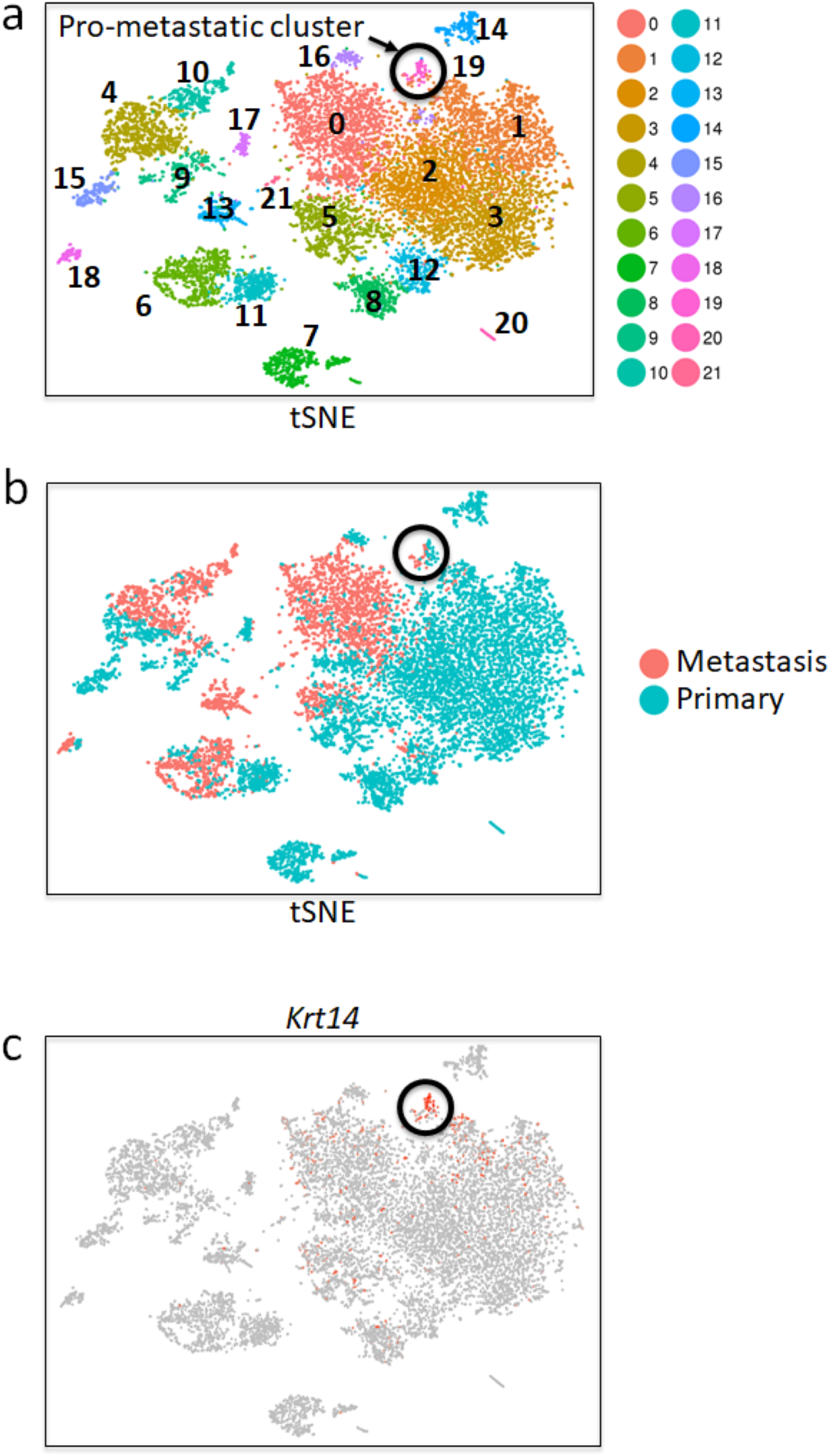
**a.** A combined tSNE plot of 7,791 primary breast tumor cells (N = 5) and 3,979 metastatic lung tumor cells (N = 3). Cluster 19 - the pro-metastatic cluster is circled. **b.** tSNE showing the cell of origin of the analysis in a. The circle shows that cluster 19 is made of both wild type (WT) primary breast cells in blue and metastatic lung (met) cells in salmon. **c.** Feature plot showing expression of *Krt14* on the tSNE localized in cluster 19.

**Supplementary Figure 7:**
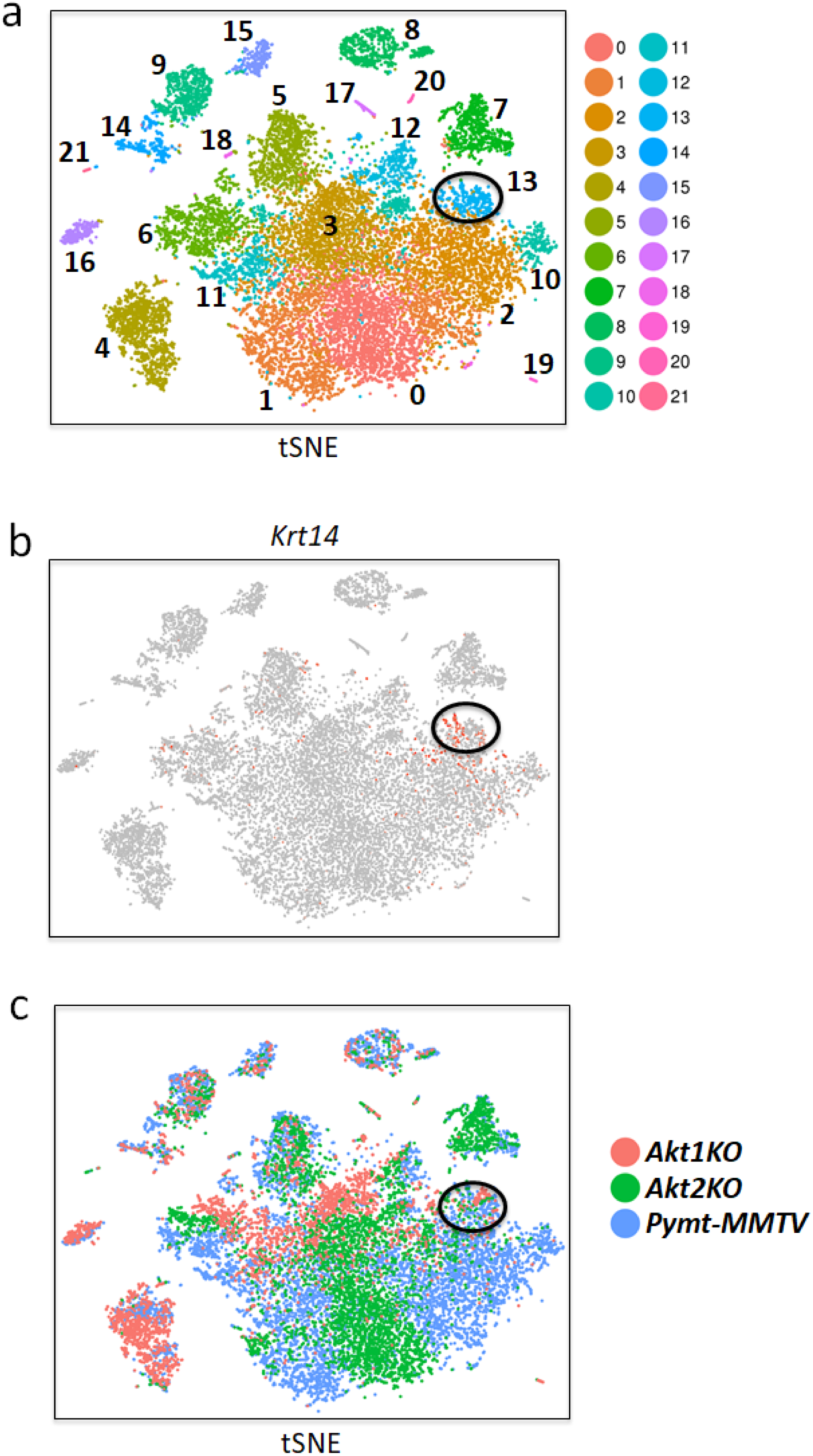
**a.** A combined tSNE plot of 7,791 primary breast tumor cells (N = 5), 3,194 cells of primary tumors following systemic *Akt1* deletion (N = 3), and 4,647 cells of primary tumors following systemic *Akt2* deletion (N = 3). Cluster 13 - the pro-metastatic population, is circled. **b.** Feature plot showing the expression of *Krt14* on the tSNE localized in cluster 13. **c.** tSNE showing the cell of origin of the analysis in a. The circle shows that cluster 13 consists of WT cells in blue, systemic *Akt1* deletion in salmon, and systemic *Akt2* deletion in green.

**Supplementary Figure 8:**
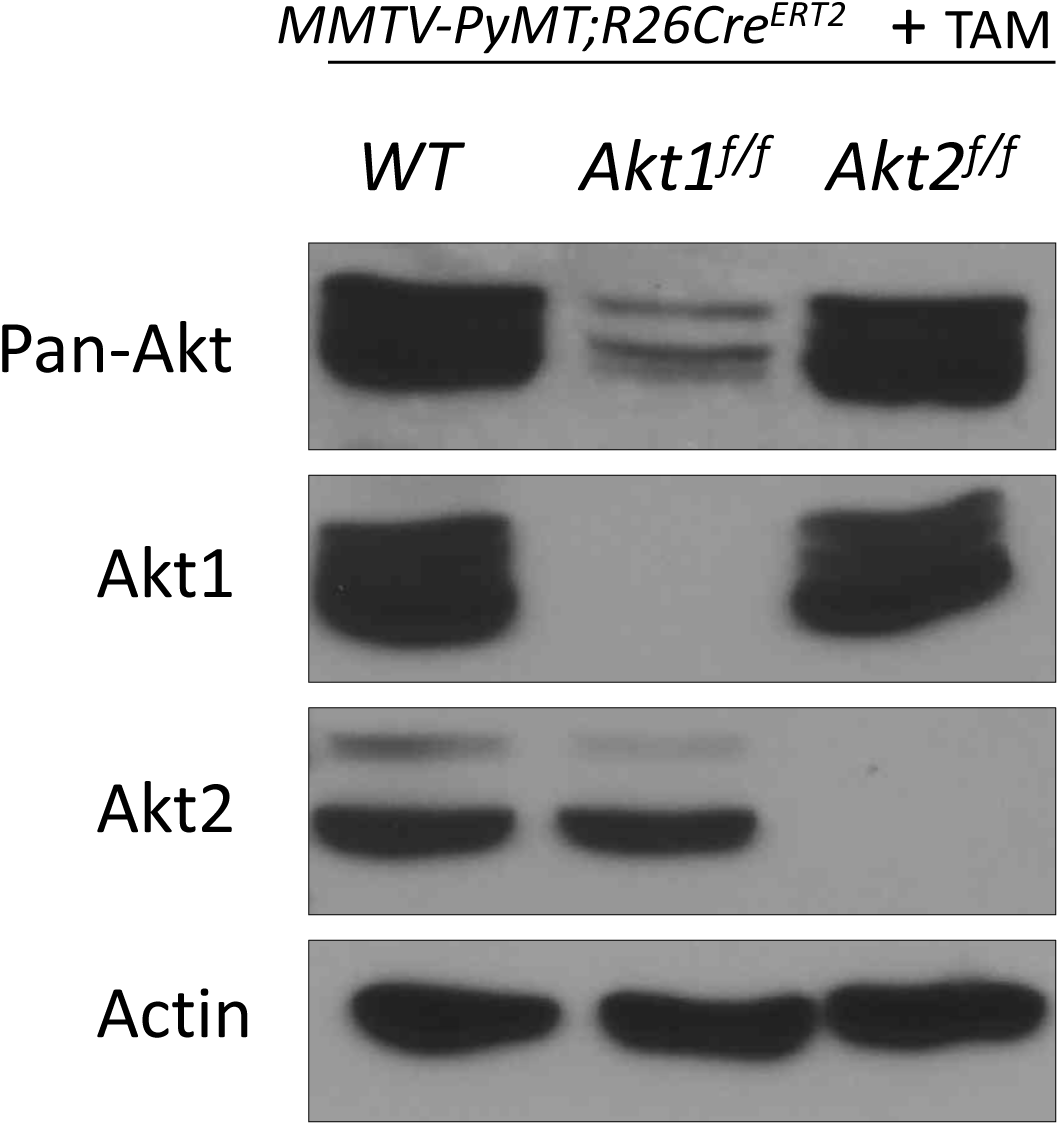
Immunoblot showing that total Akt expression (pan-Akt) in neutrophils isolated from control (WT) and systemically deleted Akt1 or Akt2 tumor-bearing mice.

